# Interferon regulates stem cell function at all ages by orchestrating mTOR and cell cycle

**DOI:** 10.1101/2022.02.03.478954

**Authors:** Damian Carvajal Ibañez, Maxim Skabkin, Jooa Hooli, Santiago Cerrizuela, Manuel Göpferich, Adrien Jolly, Marc Zumwinkel, Matilde Bertolini, Thomas Höfer, Guenter Kramer, Simon Anders, Aurelio Telemann, Anna Marciniak-Czochra, Ana Martin-Villalba

## Abstract

Stem cells show intrinsic interferon signalling, which protects them from viral infections at all ages. In the ageing brain, interferon signalling in stem cells also reduces their ability to activate. Whether these functions are linked and at what time interferons start taking on a role in stem cell functioning is unknown. Additionally, the molecular link between interferons and activation in neural stem cells and how this relates to productivity is not well understood. Here we combine single-cell transcriptomics, RiboSeq and animal models of interferon to show that this pathway is important for proper stem cell function at all ages. Interferon orchestrates cell cycle and mTOR activity to post-transcriptionally repress Sox2 and drive the exit from stem cell activation. The interferon response then decreases in the subsequent maturation states. Mathematical simulations indicate that this regulation is beneficial for the young and harmful for the old brain. Our study establishes molecular mechanisms of interferon in stem cells and interferons as genuine regulators of stem cell homeostasis and a potential therapeutic target to repair the ageing brain.

## Introduction

In the adult brain, stem cells residing in the ventricular-subventicular zone (vSVZ) generate olfactory bulb interneurons that are crucial for fine-tuning odour discrimination. For neuronal production, neural stem cells (NSCs) transit from a dormant to an activation state to produce transient amplifying progenitors (TAPs) and finally neuroblast (Urbán *et al*, 2019). These neuroblasts migrate along the rostral migratory stream toward the olfactory bulb (OB), where they mature into olfactory bulb interneurons. As the animal age activation of NSCs decreases, while interferon signalling increases (Kalamakis *et al*, 2019; Baruch *et al*, 2014). This age-related interferon response is highest in the neighbouring cells but also visible in NSCs (Kalamakis *et al*, 2019). Apart from their function in homeostasis, NSCs become activated upon injury to produce neurons and other glia cells (Delgado *et al*, 2021). This injury response is in part mediated by interferons (Llorens-Bobadilla *et al*, 2015; Kyritsis *et al*, 2012). Interferons (IFNs) are cytokines known to modulate the innate and adaptive immune response upon infections and injury (Mazewski *et al*, 2020). The interferon family is composed of type I, II and III IFNs. While type I and II IFNs are sensed ubiquitously in the body, type III IFN response is restricted to immune and epithelial cells. Type I and II IFNs activate the canonical JAK/STAT signalling pathway through IFN receptor α (IFNAR) and γ (IFNGR), respectively, leading to the transcription of a subset of interferon stimulated genes (ISGs) (Stanifer *et al*, 2020; Alspach *et al*, 2019). Despite the ubiquitous expression of IFNAR and IFNGR in stem cells, they show an attenuated response to IFN compared to differentiated counterparts (Wu *et al*, 2018). Instead, stem cells, including neural stem cells (NSCs), rely on intrinsic expression of ISGs to prevent viral infection (Wu *et al*, 2018). Whether this intrinsic interferon signalling that is observed in NSCs already in young animals regulate stem cell function has not been addressed. In addition, whether the age-related increased in interferon response is independent of the intrinsic interferon response is similarly unexplored. Interestingly, recent ageing studies hinted at a faint basal interferon response already in the young homeostatic brain, albeit they were technically unable to characterise it (Kalamakis *et al*, 2019). Moreover, previous studies focused only on transcriptional control of ISGs while the post-transcriptional regulation of stemness factors (Baser *et al*, 2019) upon IFN exposure in NSCs remains elusive. Understanding how the positive and negative functions of interferon are molecularly wired in stem cells in the young and old brain is mandatory to provide regenerative therapy for a better ageing.

## Results

### Interferon regulates neural stem cells in the young and ageing brain

IFNs are known regulators of NSCs reaction to injury and infection. To characterise a potential role of IFNs in NSCs homeostasis, we first examined the individual transcriptomes of NSC and their progeny in mice lacking the type-I (IFNA) and -II (IFNG) interferon receptors (IFNAGR^KO^) and in wild-type mice (IFNAGR^WT^). To capture changes in NSCs dynamics across ages we examined young (2-3 months old) and old mice (17-24 months old). To this end, we profiled the transcriptomes of 15,548 individual NSCs, their progeny (Tlx-mediated eYFP+ cells) and neighbouring microglia and endothelial cells isolated from vSVZ, the RMS and the olfactory bulb (Fig 1A and EV1A). Using previously defined scRNAseq markers (Kalamakis *et al*, 2019; Llorens-Bobadilla *et al*, 2015) we identified dormant NSC (qNSC1), primed-quiescent NSCs (qNSC2), active NSC (aNSC), TAPs and neuroblasts (NBs) (Fig 1B and EV1B). Next, we aimed at assessing the strength of the interferon signalling in cells along the different transitions into neuronal differentiation. Previous single-cell analysis on NSCs proved scRNAseq to be underpowered to categorize basal inflammatory signatures in the brain (Kalamakis *et al*, 2019). To maximize the power of our interferon response analysis, we explored the specific NSC type-I IFN response by treating NSCs with IFN-β *ex-vivo*. First, applying Cycleflow (prepint: Jolly *et al*, 2020) to evaluate cell cycle progression, we show that IFN-β arrested NSCs in the G_0_ quiescent state *ex-vivo* (Fig 1C and EV2), mimicking the effect of IFN-increased quiescence as already suggested for the old brain (Kalamakis *et al*, 2019). We thereafter addressed the molecular response at transcriptional and post-transcriptional level of NSCs to IFN-β *ex-vivo* via Ribo-Seq (Fig 1D). Analysis of the transcriptional response identified a strong upregulation of ISGs in NSCs (Fig 1E), opposed to the suggested attenuated capacity of stem cells to build interferon responses (Wu *et al*, 2018). The 300 highest-expressed genes (Fig 1E) were used to generate a NSC-specific type-I IFN response signature further used for detection of this response in our single cell NSC-lineage from IFNAGR^WT^ and IFNAGR^KO^ young and old vSVZ.

**Figure 1.**
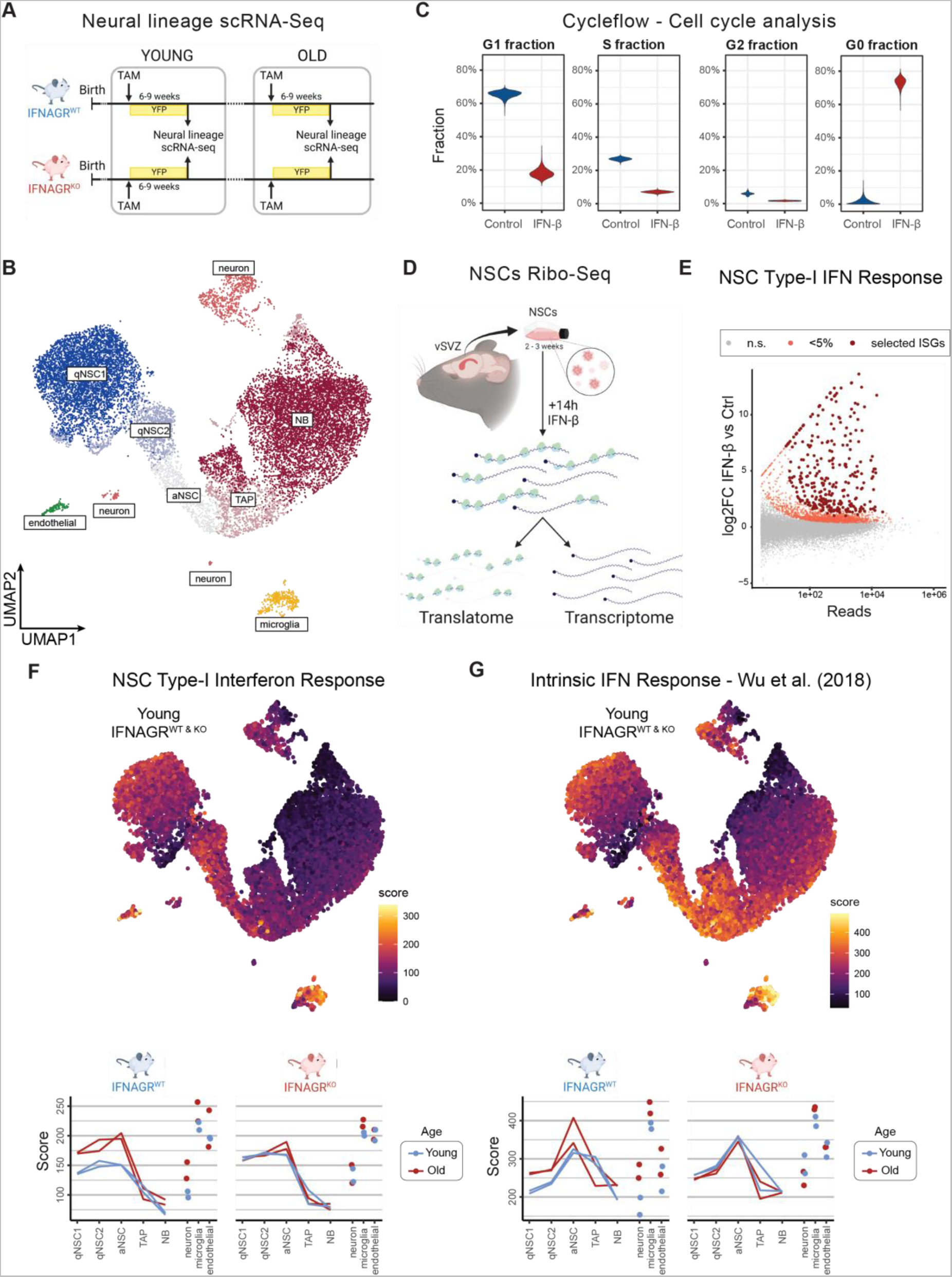
Interferon signalling regulates stem cells at all ages and decreases in neural progenitors. **A.** Experimental layout for scRNA-Seq in young and old mice lacking interferon receptors. All mice represent TiCY (Tlx reporter, see methods) that are either IFNAGR WT or KO. TAM (Tamoxifen). See also Fig EV1. **B.** UMAP embedding showing the 15,548 single cells in this analysis with their cell types. See also Fig EV1. n = 2 biological replicates per age and genotype. **C.** Cell cycle fractions inferred from Cycleflow in IFN-β-treated NSCs. See also Fig EV1. n = 3 biological replicates. **D.** Schematic representation of Ribo-Seq pipeline in NSCs. **E.** MA plot showing differential expression between IFN-β 16h treated bulk cells and Control cells. Light red dots are significantly upregulated (one-sided Wald test), dark red dots are the top 300 genes selected as the “NSC Type-I Interferon Response”. **F.** Scores computed for the NSC Type-I Interferon Response signature displayed in the UMAP embedding for young cells (with colours clipped to the range seen in the lineage cells) and averaged for the cell types in our analysis at varying ages in IFNAGR^WT^ and IFNAGR^KO^ cells. n = 2 biological replicates per age and genotype. **G.** Scores computed for the Wu et al. (2018) intrinsic interferon response gene set displayed in the UMAP embedding for young cells (with colours clipped to the range seen in the lineage cells) and averaged for the cell types in our analysis at varying ages in IFNAGR^WT^ and IFNAGR^KO^ cells. n = 2 biological replicates per age and genotype.

Scoring of the NSC type-I interferon response signature indicated that the interferon response is already present in young WT individuals and it fluctuates dynamically along the lineage (Fig 1F and EV1D-E). While intermediate progenitors (TAPs and NBs) score the lowest, stem cells and mature neurons score the highest type-I interferon response. Interestingly, stem cells and neurons are also particularly responsive in the old brain, while TAPs and NBs remain unaffected by the age-related increase of IFNs (Fig 1F, lower panel). Of note, we found expression of type-I IFN receptors in all cell types along the lineage (Fig EV1C), in addition to the recently reported expression of type-II IFN (Dulken *et al*, 2019). This underscores the relevance of interferons both in the young and the old brain and reveals neural stem cells as their preferential target to modulate neurogenesis. In agreement, stem cells lacking interferon receptors (IFNAGR^KO^) display a dysregulated type-I interferon response which remains oblivious to ageing (Fig 1F, lower panel). Strikingly, even in IFNAGR^KO^ mice, stem cells retain a higher type-I interferon response than TAPs and NBs. This supports the notion from Wu. et al. 2018 that stemness confers an intrinsic interferon response (Wu *et al*, 2018) as compared to their progeny. However, Wu. Et al. 2018 proposed stem cells to be refractory to IFNs, while we show that NSCs *in-vivo* display an interferon response that relies on IFN receptors both in the young and old brain (Fig 1G). Moreover, loss of IFN receptors increased the intrinsic levels of IFN signalling. This confirms our hypothesis that interferon modulates neural stem cells already in young adults, albeit the molecular underpinnings of such regulation remains elusive.

### IFN-β exerts a biphasic control of mRNA translation in NSCs

To describe the molecular mechanisms downstream of IFN in NSCs, we focused on the post-transcriptional changes exerted by these cytokines. IFNs can modulate mRNA translation in differentiated cells (Mazewski *et al*, 2020), a process that is of key importance in the regulation of quiescence in stem cells (Tahmasebi *et al*, 2019; Blanco *et al*, 2016; Baser *et al*, 2017, 2019). We thus hypothesized that IFN-β could also affect mRNA translation in NSCs, leading to their quiescence induction through G_0_ arrest (Fig 1C). To address this, we examined polysome profiles and Ribo-Seq of NSCs following short (2h) or long (14h) exposure to IFN-β. Analysis of the polysome profiles of IFN-β-treated NSCs revealed a mild and transient increase followed by a strong decrease of the heavy polysomal fractions, as compared to untreated controls (Fig 2A,B). Despite this IFN-β’s biphasic control of global translation, effects varied per gene (Fig 2C,D)).Gene set enrichment analysis of of translational efficiency (TE) ranked genes identified specific subsets of transcripts exhibiting either a steady enhanced or repressed translation (Fig 2C,D and EV3A,B). mRNAs encoding DNA replication and cell cycle-related proteins were steadily repressed, in agreement with the detected arrest in G_0_. Conversely, interferon stimulated genes (ISGs), crucial components of the antiviral response (Mazewski *et al*, 2020), exhibited a gradual increase of TE. This enhanced translation was confirmed by RT-qPCR of the polysomal fractions for a subset of ISGs including *Ifit3* and *Irf9* (Fig EV3C). Notably, IFN-mediated translation of ISGs had always been linked to upregulation of cap-dependent translation (Kroczynska *et al*, 2014; Mazewski *et al*, 2020). How translation of these ISGs is maintained despite a global downregulation of translation remains unclear. Overall, this shows that interferon induces a biphasic control of mRNA translation in NSCs beyond transcriptional activation of ISGs.

**Figure 2.**
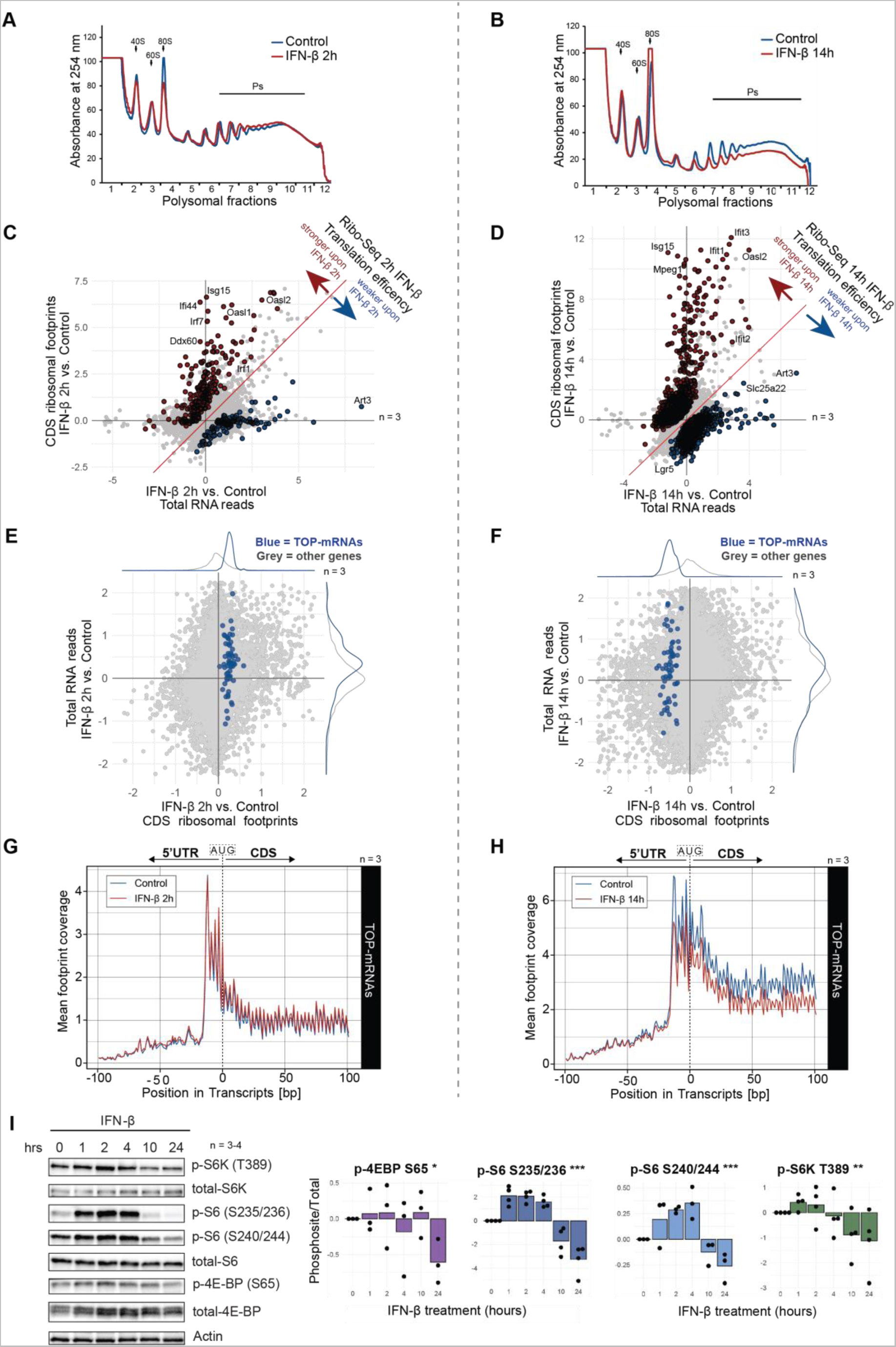
IFN-β controls mTORC1 shaping a biphasic regulation of mRNA translation in NSCs. **A,B.** Representative polysome profiles of NSCs treated with IFN-β for 2h and 14h treatment. Arrows indicate the 40S, 60S and 80S subunits of the ribosome. **C,D.** Results of Ribo-Seq depicting translation efficiency as the interaction of log2 fold changes (LFC) between footprints (ribosome protected reads) and total RNA at 2h and 14h IFN-β treatment. FDR < 10%, LR-Test in DESeq2. Genes with a p value<0.1 after FDR correction are highlighted. For associated GO terms see also Fig EV3. n = 3 biological replicates. **E,F.** Ribo-Seq results of 2h and 14h IFN-β treatments. Footprints mapped to the CDS are referred to as “CDS counts”. 5’ terminal oligopyrimidine motif-containing mRNAs (TOP-mRNAs) highlighted in blue. n = 3 biological replicates. **G,H.** Coverage profile of ribosome-protected reads (footprints) of TOP-mRNAs in 2h and 14h IFN-β treatment. Nucleotide 0 depicts the start codon (AUG) marking the interface between 5’ untranslated region (5’ UTR) and coding sequence (CDS). **I.** Representative WB and phosphorylation levels quantification (log2FC) of the mTOR-related proteins ribosomal protein S6 kinase (S6K), ribosomal protein S6 (S6) and eIF4E-binding protein (4E-BP) from NSCs treated with IFN-β and normalized to control (t=0h). Bars represent the mean value. One-way ANOVA for the biphasic response test (See Methods for details). n = 3-4 biological replicates. p ≡ p-value. *p<0.05, **p<0.01, ***p<0.001.

### mTOR contributes to the IFN-induced biphasic control of mRNA translation

To dissect the molecular drivers of the IFN-induced biphasic control of mRNA translation, we examined the implication of mTOR. mTOR is a key checkpoint in the control of metabolic pathways, including regulation of cap-dependent translation (Liu & Sabatini, 2020). Type-I, -II and -III IFNs have been reported to activate the PI3K/mTOR pathway in differentiated cell types (Lekmine *et al*, 2003, 2004; Ivashkiv & Donlin, 2014; Syedbasha *et al*, 2020). There are also recent reports on mTOR inhibition by IFN-β and IFN-γ (Su *et al*, 2015; Vigo *et al*, 2019). To assess the impact of mTOR in the observed IFN-β’s biphasic control of global translation in NSCs, we inspected the TE of mRNAs containing the 5’ terminal oligopyrimidine (TOP) motif, referred to as TOP-mRNAs (Avni *et al*, 1996). Translation of TOP-mRNAs is susceptible to mTOR activity (Thoreen *et al*, 2012), which makes it a faithful readout of mTOR activity. We observed that IFN-β controlled TOP-mRNAs in a biphasic manner, reproducing the early mild up-and late drastic down-regulation of translation (Fig 2E,F and EV3D). Further inspection of the Ribo-Seq profiles of expressed TOP-mRNAs confirmed this biphasic regulation with a differential abundance of footprints in their coding sequence (CDS) upon IFN-β treatment (Fig 2G,H). To fully confirm the implication of mTOR, we next examined the phosphorylation state of mTOR pathway components. The same biphasic regulation of phosphorylation was observed in the downstream substrates of mTOR, ribosomal protein S6 kinase (S6K), ribosomal protein S6 (S6) and eIF4E-binding protein (4E-BP) (Fig 2I). In particular, Thr389 phosphorylation of S6K, a direct target of mTORC1 (Ma & Blenis, 2009), confirmed the specific modulation of mTORC1 by IFN-β in NSCs.

### The biphasic control of mRNA translation by IFN-β involves modulation of mTORC1 and p-eIF2α

We next set out to address the molecular underpinnings of mTOR biphasic modulation. The control of mTOR by type-I IFN in different cell systems mainly relies on the crosstalk of the IFN-activated JAK/STAT pathway with the PI3K pathway (Mazewski *et al*, 2020; Platanias, 2005). In this scenario, IFN activates JAK1/TYK2 kinases that interact with the insulin receptor substrate 1 (Irs1), which modulates the activity of the PI3K-pathway components Akt and TSC1/2 (Platanias, 2005; Kim & Guan, 2019). Notably, we found that upon IFN-β treatment, phosphorylation of Akt^Thr308^ and TSC2^Thr1462^ not only increased, but followed a bimodal mode of regulation (Fig 3A), as for mTORC1 (Fig 2). This indicates that the crosstalk between IFN-β and the PI3K pathway causes the early mTORC1 upregulation. However, additional mechanisms are necessary to explain a late downregulation of the PI3K pathway upon sustained IFN-β treatment (Muendlein *et al*, 2020). To be able to follow kinase activities downstream of IFN-β in a broader manner, we acquired the proteomics and phospho-proteomics profiles of NSCs treated with IFN-β or vehicle at 2h and 14h. Analysis of these profiles revealed a downregulation of p-Akt^Ser473^, an mTORC2-dependent site (Sarbassov *et al*, 2005), only after long exposure to IFN-β (Appendix Table 1). Congruently, hyperactivation of mTORC1 can trigger negative feedback loops involving mTORC2 and Irs1/2, that reverse the initial IFN-β-mediated PI3K-Akt activation (Hsu *et al*, 2011; Rozengurt *et al*, 2014) (Fig 3B). Indeed, we also found that the early upregulation of the activity read-out of IRS1 (Esposito *et al*, 2001), p-Irs1^Tyr608^, faded upon long exposure to IFN-β. Additionally, p-Irs1^Ser1134^ and p-Irs1^Ser1137^ displayed a biphasic trend, correlating with the activation status of mTOR (Copps & White, 2012; Karlsson *et al*, 2021; Langlais *et al*, 2011) (Appendix Table 1). This suggests that prolonged IFN-β exposure in NSCs induces negative feedback loops on Akt that ultimately inhibits mTORC1 (Fig 3B). According to our results, these two temporal components of the biphasic regulation of mTOR rely on TSC1/2 modulation by Akt. To validate this, we generated heterozygous TSC2 mutant NSC clones by CRISPR/Cas9 (TSC2^mut^). NSCs where CRISPR/Cas9 did not reduce TSC2 protein levels were treated as control (TSC2^ctrl^) to exclude off-target or manipulation-derived artefacts (Fig EV4A). TSC2^mut^ NSC clones had a basally increased mTOR activity and displayed a milder inhibition of global translation in response to IFN-β (Fig EV4B,C). As hypothesized, the biphasic regulation of mTORC1 activity, as measured by p-S6K^Thr389^, was strongly blunted in TSC2^mut^ NSCs (Fig EV3D). This denotes the PI3K crosstalk and the inhibitory feedback loops converging on TSC1/2 as the main drivers of the biphasic mRNA translation control exerted by IFN-β in NSCs.

**Figure 3.**
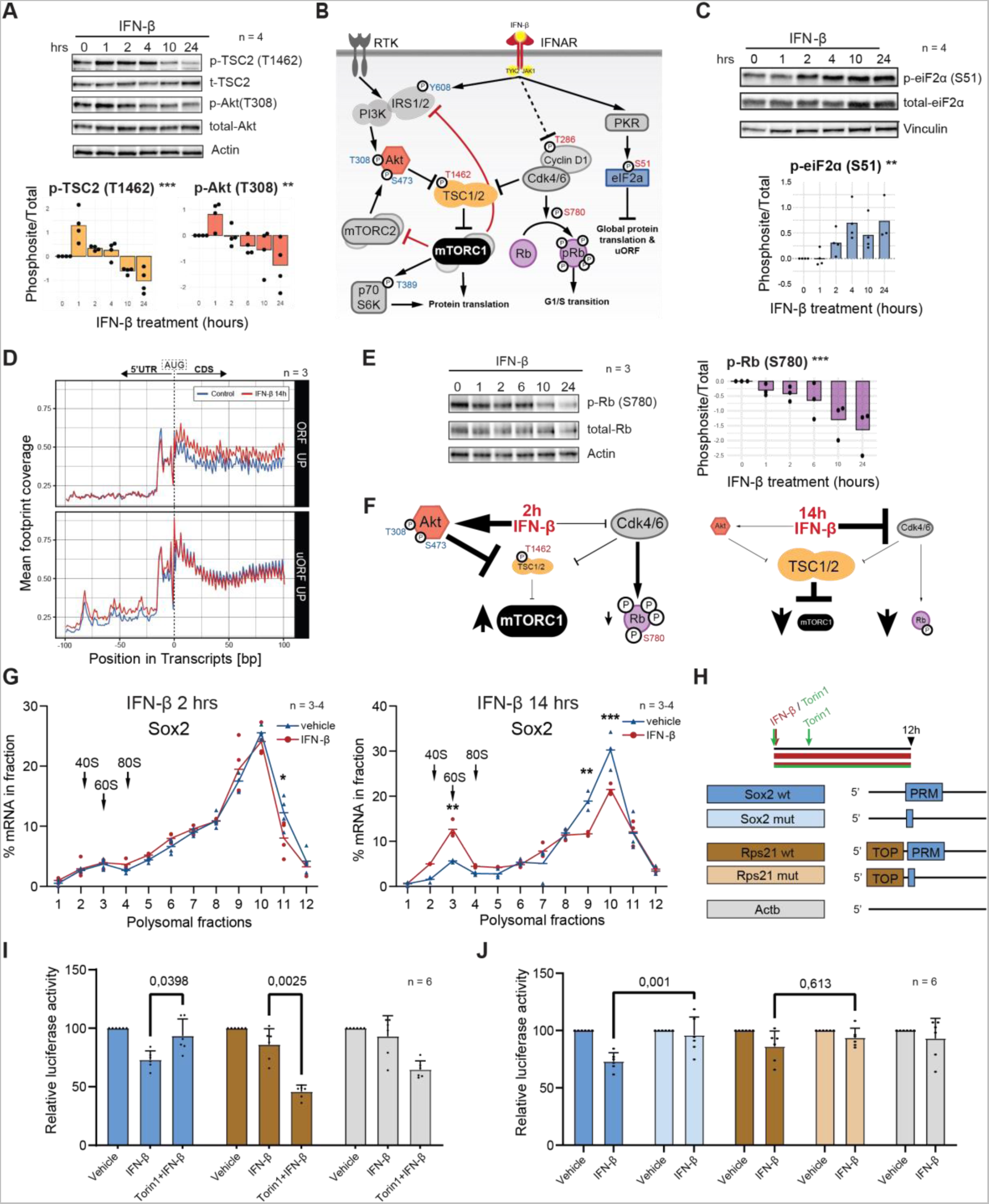
IFN-β’s biphasic control of mTORC1 uncouples growth and cell cycle and represses Sox2 translation via its 5’ PRM motif. **A.** Representative WB image and quantification (log2FC) of p-TSC2^Thr1462^ & p-Akt^Thr308^ from IFNβ-treated NSCs normalized to control (t=0h). Bars represent the mean value. One-way ANOVA for the biphasic response test (See Methods for details). n = 4 biological replicates. **B.** Schematic representation of the type-I interferon signalling pathway and its crosstalk with the receptor tyrosine kinase (RTK) pathway. Activatory (blue) and inhibitory (red) phosphosites are depicted. Red inhibitory lines depict late mTORC1-derived inhibitory feedback loops. **C.** Representative WB images and quantifications (log2FC) of p-eIF2α^S51^ in IFN-β-treated NSCs normalized to control (t=0h). See also Figure S3. Bars represent the mean value. n = 3 biological replicates. Spearman’s rank correlation test. **D.** Coverage profile of ribosome-protected reads (footprints) in genes with upregulated canonical (upper) or alternative upstream (lower) ORFs upon extended IFN-β treatment. Nucleotide 0 depicts the start codon (AUG) marking the interface between 5’ untranslated region (5’ UTR) and coding sequence (CDS). **E.** Representative WB image and quantifications (log2FC) of p-Rb1^Ser780^ from IFN-β-treated NSCs normalized to control (t=0h). Bars represent the mean value. n = 3 biological replicates. Spearman’s rank correlation test. **F.** Schematic representation of the prevailing signalling pathways driving the early inhibition and late activation of mTORC1. **G.** Polysome profiling (RT-qPCR) of Sox2 mRNA upon 2h and 14h IFN-β treatment. Hyphens represent mean of biological replicates. Arrows represent the 40S, 60S and 80S subunits of the ribosome. n = 3-4 biological replicates. Two-way ANOVA with Šídák’s multiple comparison test was computed. **H.** Experimental layout of IFN and Torin1 treatments (upper panel). Schematic representation (lower panel) of the 5’UTR constructs priming the *renilla* luciferase. TOP = 5’Terminal Oligopyrimidine motif; PRM = 5’ Pyrimidine Rich Motif. **I,J.** Luciferase activity controlled by the upstream 5’UTR fragment from *Sox2* (WT and mutant), *Rps21* (WT and mutant) and *Actb*. Data is normalized to vehicle and is represented as mean ± SD. n = 6 biological replicates. Two-way ANOVA with Tukey’s multiple comparisons test (p-value specified). *p<0.05, **p<0.01, ***p<0.001.

Along with the inhibitory feedback loops, transcriptionally upregulated ISGs, such as PKR, promote a delayed inhibition of global mRNA translation (Ivashkiv & Donlin, 2014). PKR (*Eif2ak2*) is activated by dsRNA and phosphorylates the translation factor eIF2α at Ser51, resulting in a global inhibition of mRNA translation initiation (Gal-Ben-Ari *et al*, 2019). Certainly, NSCs exposed to IFN-β induced an increase in p-eIF2α^Ser51^, which intensifies the inhibition of mRNA translation already exerted by late inhibition of mTORC1 (Fig 3C). Of note, extended exposure to IFN-β also increased PKR protein levels and phosphorylation in poorly-described residues (Piazzi *et al*, 2020) (Fig EV4E; Appendix Table 1), suggesting a late PKR activation. In addition, activated PKR can inhibit Irs1 (Nakamura *et al*, 2010), contributing to the late mTORC1 feedback inhibitory loops. This shows how the late phosphorylation of eIF2α upon sustained IFN-β exposure contributes to the biphasic control of mRNA translation and suggests PKR as its upstream regulator.

In addition to the regulation of global mRNA translation, p-eIF2α^Ser51^ is associated with the alternative use of upstream open reading frames (uORFs) located in the 5’ untranslated region (5’UTR) (Young & Wek, 2016). IFN-β increased local 5’UTR footprint density in a subset of genes while maintaining unaltered levels in the CDS, indicating alternative uORF usage (Fig 3D). Consistently, while translation upregulation in the CDS shifted mRNAs to heavy polysomes, a higher ribosomal density in uORFs shifted the associated mRNAs into light polysomes (Fig EV4F).

### IFN-β transiently uncouples mTOR and cell cycle in NSCs

It did not remain unnoticed that the transient increase of mRNA translation induced by IFN coincides with the inhibition of cell cycle in NSCs (Fig 1C). We therefore assessed how the type-I IFN response uncouples translation and cell cycle in NSCs. Given that NSCs remained significantly longer in G_1_ upon IFN-β treatment (Fig EV2B), we hypothesized that IFN could trigger the inhibition of the G_1_/S checkpoint regulators Cdk4/6. Cdk4/6 have recently emerged as a dual regulator of cell growth and proliferation via TSC2, and together with CyclinD1 they modulate proliferation of NSCs and neurogenic progenitors in the adult brain (Romero-Pozuelo *et al*, 2020; Lange *et al*, 2009; Artegiani *et al*, 2011). Cdk4/6 control the G1/S transition by phosphorylation of Rb1 and consequent expression of S-phase genes (Topacio *et al*, 2019). We found that IFN-β reduced p-Rb^Ser780^, a Cdk4/6-dependent phosphosite, in a progressive manner in NSCs *ex-vivo* (Fig 3E). This confirms that NSCs transiently uncouple mRNA translation and cell proliferation upon exposure to IFN-β. Interestingly, IFN-β treatment also led to the phosphorylation of CyclinD1 at Thr286 (Appendix Table 1). This phosphosite is involved in cytoplasmic translocation and degradation of CyclinD1 (Guo *et al*, 2005), presenting cyclinD1 as a direct target of the IFN response in NSCs.

Overall, our data provide molecular insights of the biphasic control of mTOR by IFNs in stem cells. We show that an early activation of PI3K-Akt masks a gradual inhibition of Cdk4/6 that only gains relevance upon extended IFN-β incubation and complements the late inhibitory feedback loops shutting down mTOR (Fig 3F).

### Dual control of mTOR and cell cycle by IFN-β represses Sox2 translation via its 5’ pyrimidine rich motif

A dual regulation of mTOR and cell cycle is needed to repress translation of *Sox2* and allow exit of an activated stem cell state (Baser *et al*, 2019). However, homeostatic cues orchestrating this dual regulation in NSCs are yet missing. We therefore wondered whether type-I IFN, via its dual effect on TOR and cell cycle, would be one upstream post-transcriptional regulator of *Sox2* expression. To test this we first examine expression of Sox2 in polysomal profiles obtained from untreated and IFN-β-treated NSCs. A*s opposed to TOP-mRNAs, Sox2* remained oblivious to the initial increase of mTORC1 activity and even exhibited a subtle repression at 2h after IFN-β (Fig 3G, left panel, and EV3D). Notably, *Sox2* translation was strongly repressed in NSCs following long exposure to IFN-β (Fig 3G, right panel). Since IFN-β-mediated repression was much stronger than the one we previously measured following TOR inhibition and cell cycle arrest (Baser *et al*, 2019), we wondered whether the early decoupling of mTOR and cell cycle exerted by IFN-β would be required to more effectively repress *Sox2* translation. In order to evaluate this hypothesis, we blocked the early mTOR activation with Torin1 in IFN-β-treated NSCs (Fig 3H). To simultaneously examine the involvement of the 5’UTR of *Sox2* in this repression, we established a luciferase assay in which we cloned the 5’UTR region of *Sox2* as well as of the TOP-containing mRNA *Rps21* upstream of the *Renilla* luciferase CDS (Fig 3H). Sole mTOR inhibition by Torin1 treatment effectively repressed translation of the TOP-mRNA *Rps21* but failed to repress *Sox2* (Fig EV4G), in agreement with previous findings (Baser *et al*, 2019). Conversely, dual control of mTOR and cell cycle by IFN-β treatment effectively repressed Sox2 (Fig 3I). However, IFN-β repression of *Sox2* was prevented by Torin1 (Fig 3I). This underscores the importance of IFN-β as a *bona-fide* regulator of Sox2 translation by uncoupling of cell cycle and mTOR activity in NSCs.

In addition, we recently proposed that a pyrimidine rich motif (PRM) present in the 5’ UTR of relevant stem genes could be responsible for post-transcriptional repression of factors involved in stem cell exit (Baser *et al*, 2019). PRM is present in different mRNAs downregulated at the onset of differentiation such as *Sox2* and *Pax6* and can co-localize with the TOP motif as in the case of *Rps21* (Baser *et al*, 2019). To evaluate the contribution of the PRM to the repression of *Sox2*, we mutated the PRM using *Rps21* as control (Fig 3H). The deletion of the PRM attenuated the repression of *Sox2* exerted by IFN-β in NSCs (Fig 3J) and did not affect expression of the PRM-mutated *Rps21*, driven by the prevalent TOP motif. This highlights the role of the 5’UTR PRM motif in regulating translation of pioneer factors such as Sox2 in stem cells.

### Interferon regulates stem cell function and fine-tunes progenitor production across all ages

Altogether we have shown that IFN targets NSCs in the young and old brain and that IFN blocks stem cell activation through a biphasic control of mRNA translation and repression of Sox2. In addition, we show that the interferon response of NSCs in the natural environment of the brain relies on external interferons both in young and aged individuals (Fig 1F). In order to examine how IFNs fine-tune the adult vSVZ’s neurogenesis across the animal’s life-span, we examined NSC dynamics in young and old IFNAGR KO mice. To this end, we quantified the pool of NSCs (Prom1^+^GLAST^+^) within the vSVZ via flow cytometry and compared them to wild-type (WT) controls. In addition, we used the fraction of active NSCs, i.e., the number of cycling cells among the BrdU-retaining cells, previously acquired in WT and IFNAGR KO animals^3^. These data were analyzed using a mathematical model previously established and validated in wild-type animals (Kalamakis *et al*, 2019). This model describes the dynamics of quiescent, active and neurogenic progenitor populations and allows linking those to age-dependent changes in the cell parameters: (i) activation rate and (ii) self-renewal probability (Fig 4A, Appendix Data 1). For wild-type data, as already reported (Kalamakis *et al*, 2019), it pointed to a profound reduction of the NSC activation rate and a slight increase in self-renewal probability as the parameters responsible for a deceleration in NSC decline in old mice. Notably, this deceleration is needed to maintain a minimal amount of NSCs that otherwise would be fully depleted at older ages. In IFNAGR KO, NSC depletion exhibited faster dynamics and yet stabilized at counts of around 150 cells per mouse, as in WT mice (Fig 4B). The mathematical model applied now to the IFNAGR KO data predicted a constant activation rate of NSCs and an earlier time-dependent regulation of the probability of NSC self-renewal (Fig 1C). The latter explained the observed saturation in the NSC decline. This unveils self-renewal as a key second layer of regulation, following modulation of the activation rate, which had been previously neglected (Kalamakis *et al*, 2019).

**Figure 4.**
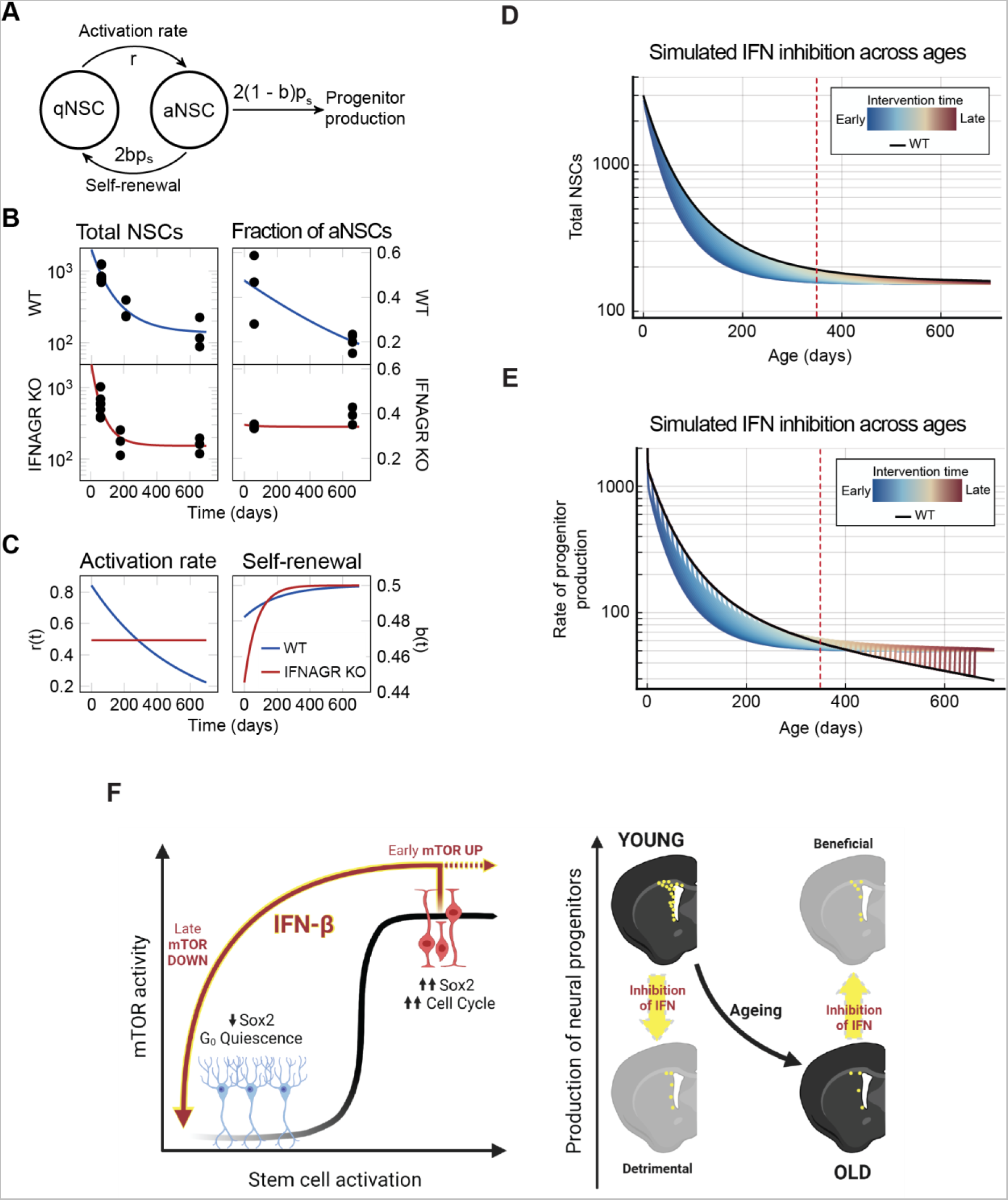
Interferon regulates stemness and fine-tunes progenitor production at all ages. **A.** Graphical depiction of the mathematical model describing the activation (at rate r) of quiescent neural stem cells (qNSC) into active NSC (aNSC) followed by their progression through cell cycle (at rate p_s_) into either two downstream progenitors (at probability 1−b) or two NSCs (at probability b) that return to quiescence. In models with time-dependent parameters, *b*≡*b*(*t*) and *r*≡*r*(*t*), See Appendix Data 1 for details. **B.** vSVZ niche FACS quantifications of total NSCs (Total NSCs) & immunofluorescence quantification of aNSCs (Fraction of aNSCs) in WT or IFNAGR KO mice at 2, 7 and 22 months old (MO). Data from individual mice is plotted. Inferred dynamics are depicted as a blue (WT) or red (IFNAGR KO) line over time. n = 3-8 biological replicates. **C.** Inferred parameters are depicted as a blue (WT) or red (IFNAGR KO) line over time. **D.** Induced interferon knockout simulations (colored, each color represents a different intervention timepoint) against wildtype simulations (black) showing the total number of stem cells across age. Dashed red line denotes age 350 days. **E.** The rate at which progenitors are produced (2*p_s_* (1 – *b*) *aNSC*) from NSCs for induced interferon knockout simulations with interferon dependent self-renewal (coloured, each colour represents a different intervention timepoint) and WT simulations (black). Dashed red line denotes age 350 days. **F.** Synopsis abstract

Last, given the relevance of extrinsic IFNs for NSCs in youth and ageing (Fig 1), we further explored its potential role as a therapeutic target. In our models, we know that the activation rate of NSCs directly influences the production of progenitors (Appendix Data 1). Young IFNAGR KO mice have a lower activation rate of NSCs than WT, suggesting a decreased productivity. Older IFNAGR KO mice however have a higher activation rate than WT, suggesting increased productivity (Fig 4C). This response suggests a possibility for improving progenitor production in WT animals by selecting an optimal time point to block IFN function. To investigate this, we simulated a modified model with IFN-dependent switches in model parameters at arbitrary time points (Appendix Data 1). The intervention was modelled by switching the activation rate and self-renewal parameters estimated from WT to the parameters estimated for IFNAGR KO mice at a given intervention time point. This best mimic an acute clinical intervention for systemic neutralization of interferons. Interventions simulated at young ages resulted in a significant loss of NSCs, dropping from thousands to hundreds in the short term (Fig 4D). However, this effect diminished with time resulting in similar cell counts in old age (Fig 4D and EV5A). Since total NSCs are not necessarily indicative of proper functioning, we also computed the cell flux from aNSC to progenitors (progenitor production). Simulations of young age interventions showed drastically reduced progenitor production on a short time scale, which ultimately changed to an increased progenitor production in old ages (Fig 4E). Strikingly, interventions in older mice (after ∼300 days) showed to always be beneficial. We next compared the summary effect of an intervention in terms of the life-long progenitor production. The interventions at young ages had diminished progenitor production when compared to WT, while the maximal benefit of the life-long production was achieved by intervention at about 350 days of age (Fig 4E and EV5B). Curiously, this was also the age at which the activation rate for WT dropped below the constant IFNAGR KO activation rate (Fig 4C). Overall, this shows that there may be benefits to progenitor production from blocking interferon response at increasing ages, while it may have strong adverse effects on short-term productivity and stem cell counts when intervening too early. In agreement with our predictions, local blockade of CXCL10 signalling in the aged brain as well as neutralization of IFN (Baruch *et al*, 2014) increased progenitor production from vSVZ-NSCs (Blank *et al*, 2016; Kalamakis *et al*, 2019). CXCL10 is induced in endothelial cells upon systemic IFN-β, which itself is not able to cross the blood brain barrier (Blank & Prinz, 2017). Altogether, our predictions show that only a late intervention blocking IFN signalling is beneficial and unveils the relevance of homeostatic interferons for a life-long healthy adult neurogenesis.

## Discussion

### Interferons are bona fide regulators of stem cell homeostasis at all ages and potential candidates to repair the ageing brain

Stem cells exhibit an intrinsic IFN signature, i.e., constitutive expression of ISGs independent of interferon receptor stimulation, which is unique to stem cells and protects them against viral infections (Wu *et al*, 2018). In addition, Wu et al. also reported that stem cells are refractory to extrinsic IFNs. Our study shows that in the natural environment of the brain, type-I interferon ISGs are highest expressed in NSCs but also present, albeit to a lower extent, in fully differentiated new-born neurons. This IFN signature in neural stem cells relies on expression of IFN receptors both in the old and the young brain and modulates neural progenitor production along the lifespan of the individual. Our data also reveals that neural stem cells are the preferred target and indeed responsive to IFNs both *in-* and *ex-vivo*, as it is the case for their fully differentiated counterparts. It is however the intermediate maturation stages that shutdown expression of these ISGs. As opposed to previous reports claiming that the ability to respond to IFN ligands is acquired upon exit of stemness (Wu *et al*, 2018), our single cell transcriptome data show that intermediate progenitors are not responding to interferons neither in the young nor in the ageing brain. Thus, intermediate progenitor stages are refractory to extrinsic interferon and probably represent the most sensitive cells to viral infections along the neural lineage. We believe that during brain development the high number of intermediate progenitors could contribute to viral infections being a life-threatening risk at this age. In addition, our models reveal the blockade of external IFN signalling to be harmful for the young vSVZ albeit beneficial in ageing. This is supported by previous reports blocking IFN signalling in ageing (Kalamakis *et al*, 2019; Baruch *et al*, 2014) despite the recently reported IFN memory in stem cells (Haas *et al*, 2017). Altogether, our combined single-cell transcriptomics and mathematical modelling underscore the relevance of defining a critical time of intervention and unveils a previously neglected role of interferon signalling in the homeostasis of the young adult brain.

### Type-I IFN induces stem cell quiescence by a biphasic control of mRNA translation and cell cycle exit

IFNs modulate stem cells across different tissues but the molecular underpinnings of this regulations are yet poorly described (Demerdash *et al*, 2021; Zhang *et al*, 2020; Hou *et al*, 2021; Kalamakis *et al*, 2019). Performing polysome profiles and ribosome footprinting on IFN-β treated NSCs we find that IFN-β induces a biphasic control of mRNA translation. This regulation involves a transient activation of mTORC1 followed by its inhibition, as opposed to the previously reported unidirectional modulation of mTOR by type-I IFN (Mazewski *et al*, 2020). Of note, recent studies highlighted the importance of a precise pharmacological regulation of mTOR activity in NSCs, since hyperactive mTOR is associated with Alzheimeŕs Disease both in patients and mouse models (Nicaise *et al*, 2020). Future studies should address the potential role of this biphasic regulation of mTOR in brain repair. We further show that the transient activation of mTOR is the result of the crosstalk between IFN-β signalling and PI3K. This transient activation of mTOR triggers inhibitory feedback loops acting on Akt that, together with a delayed activation of PKR and phosphorylation of eIF2α, ultimately lead to the late and profound inhibition of mRNA translation. Whether this biphasic response is a hallmark of stem cells beyond the brain will be subject for future investigations.

Additionally, our data suggests that IFN-β treatment in NSCs not only induces the expression but also activates PKR in a dsRNA-independent manner. This is an undescribed feature of IFNs in stem cells as a similar response was only proposed in an interferon-selected carcinoma line harbouring oncogenic mutations (Su *et al*, 2007). Interestingly, high levels of p-eIF2α are also associated to self-renewal and quiescence induction in embryonic and adult stem cells (Friend *et al*, 2015; Zismanov *et al*, 2016). As a whole, these pathways converge in a profound shut down of translation that keeps NSCs dormant. Similarly, chronic type-I IFN promotes quiescence after a transient activation of hematopoietic stem cells (HSCs) (Pietras *et al*, 2014). Future studies should address whether this transient activation of HSCs by IFN is also driven by a biphasic modulation of mTOR.

Supporting the induction of dormancy, we find that IFN-β shifts NSCs to a quiescent G_0_ state, as opposed to the previously reported increase in cell-cycle length in quiescent NSCs (Daynac *et al*, 2016). Remarkably, inhibition of Cdk4/6 upon IFN-β treatment transiently uncouples mTOR and cell cycle in NSCs. A similar uncoupling of proliferation and protein synthesis was observed upon differentiation of skin stem cells, but the drivers of this response remained elusive (Blanco *et al*, 2016). Of note, senescent cells, a hallmark of ageing (Fulop *et al*, 2018; Zhu *et al*, 2021), steadily uncouple mRNA translation and cell cycle (Payea *et al*, 2021). Differently to NSCs, a late coupling of translation and cell cycle has not been described in senescence. Intriguingly, IFN-β is one of the main drivers of senescence. Cytosolic chromatin fragments present in senescent cells activate cyclic GMP-AMP synthase (cGAS) that, in turn, triggers the production of inflammatory factors including IFN-β, thereby inducing paracrine senescence (Glück & Ablasser, 2019). Although the prevalence of senescent cells increases with age in the neurogenic niches in the brain (Fernández-Fernández *et al*, 2012; Molofsky *et al*, 2006; Ogrodnik *et al*, 2019; Jin *et al*, 2021), how NSCs are protected from senescence remains unknown. Overall, our study describes the molecular underpinnings of interferon signalling in NSCs and reveals previously undescribed features of the IFN-β response in stem cells.

### Type-I IFN induces a biphasic control of mTOR that represses *Sox2* translation in NSCs

We recently reported that a dual regulation of mTOR and cell cycle is needed to repress translation of *Sox2* in NSCs (Baser *et al*, 2019). We now identify type-I IFN as a regulator of this response. Furthermore, we narrow down the nature of this repression to the transient activation of TOR and the presence of the recently described pyrimidine-rich motif (PRM) (Baser *et al*, 2019) present on the 5’ UTR of *Sox2*. In this recent study, we addressed the role of repression of *Sox2* translation in the transition of an activated stem cell into a differentiated neuroblast (Baser *et al*, 2019). Sox2 levels are however likewise reduced in quiescent NSCs to avoid replicative stress (Marqués-Torrejón *et al*, 2013). The determinants of directionality into a differentiated progenitor or a quiescent stem cell state following repression of Sox2 remain subject of future studies. Notably, Sox2 expression is reduced in the ageing brain, correlating with increased quiescence in NSCs (Carrasco-Garcia *et al*, 2019).

Collectively, our findings unveil interferons as *bona fide* regulators of stem cell function during homeostasis as well as in ageing. Mechanistically, IFN-β arrests NSCs at G_0_ of the cell cycle and drives a transient TOR activation followed by a profound decrease of TOR activity and global protein synthesis. This biphasic regulation is orchestrated by mTORC1, PKR and p-eIF2α. In addition, we show that the transient increase of TOR activity is required for post-transcriptional repression of PRM-containing transcripts, such as *Sox2*, that contribute to the exit from the activation state in stem cells (Fig 4F). Future studies should unveil the potential uniqueness of this biphasic response in stem cells across different tissues and its relevance for the prevention of senescence in stem cells. Our study indicates that IFN’s control of the activation rate and self-renewal of NSCs adapts the output of differentiated progenitors to demand. In young brains, with high demand, interferon would increase the output, and vice versa, decrease the number of generated progenitors in the less active ageing brain. Therefore, while the loss of IFN signalling is beneficial in the old brain, it is detrimental in young individuals. The fact that the intrinsic expression of ISGs required as antiviral defence even slightly increases after blocking interferon receptors, discloses IFNs as potential therapeutic target to improve stem cell homeostasis and repair in the brain without compromising antiviral response. However, intervention should be timed only after reaching advanced age.

## Materials and Methods

### Mice

C57BL/6 male mice were purchased from Janvier or bred in-house at the DKFZ Center for Preclinical Research. Male IFNAGRKO (Huang *et al*, 1993; Müller *et al*, 1994) (IFNAR^-/-^IFNGR^-/-^) were backcrossed to a C57BL6 background. Male TiCY mice (Tlx-CreERT2-YFP) (Liu *et al*, 2008) (TiCY [B6-Tg(Nr2e1-Cre/ERT2)1GscGt(ROSA)26Sortm1(EYFP)CosFastm1Cgn/Amv]) were crossed with IFNAGRKO mice to generate TiCY-IFNAGRKO mice ([B6-Tg(Nr2e1-Cre/ERT2)1GscGt(ROSA)26Sortm1(EYFP)CosFastm1CgnIfnar1tm1AgtIfngr1tmAgt/Amv]). Animals were housed under standard conditions and fed ad libitum. All procedures were approved by the Regierungpräsidium Karlsruhe.

### Cell culture and treatments

For NSC isolation, the subventricular zones from 8-12 weeks old male C57BL/6 mice were isolated as described (Mirzadeh *et al*, 2010). The tissue was dissociated using the Neural Tissue Dissociation Kit with Papain (Miltenyi Biotec) following the manufacturer’s instructions. Cells were cultured in Neurobasal A Medium supplemented with 2% B27 Supplement serum-free, 1% L-Glutamine (all from ThermoFisher), 2 µg/ml of heparin, 20 ng/ml of human basic FGF (ReliaTech) and 20 ng/ml of human EGF (Promokine). Cells were maintained in a 37°C, 5% CO_2_ incubator.

For the treatment with interferon, mouse recombinant IFN-β (Millipore) diluted in DPBS with 0.1% Bovine serum albumin BSA (Roche) were added directly to the culture media at final concentration of 40 u/ml for the indicated time. Control cells were treated with the same volume of 0.1% BSA in PBS.

For the treatment with Torin1 (Cayman Chemical), Torin1 was dissolved in 100% DMSO and added to cells at 250 nM for the indicated time. Control cells were treated with the same volume of DMSO.

### CRISPR-mediated generation of TSC2^+/-^ NSCs

CRISPR-Cas9 gRNAs targeting exon 2 of *Tsc2* (ENSMUSG00000002496) were designed using the CCTop target online predictor (Stemmer *et al*, 2015) and cutting efficiency was evaluated by T7EI Assay using T7 Endonuclease I (NEB). We selected the TSC2Ex2-gRNA (TGTTGGGATTGGGAACATCGAGG) and cloned it into the Cas9-containing plasmid pSpCas9(BB)-2A-GFP (Addgene) as described (Ran *et al*, 2013) (see Appendix Table 2).

For nucleofection, 1 x 10^6^ NSCs were mixed with 2.5 µg of corresponding DNA using Amaxa P4 Primary Cell 4D-Nucleofector X Kit S (Lonza) with CA137 programme in a 4D-Nucleofector X Unit (Lonza). Cells were incubated for 24 hours at 37°C and 5% CO_2_. GFP-positive single NSCs were FACS-sorted into 96-well plates. Single NSC colonies were expanded and TSC2 abundancy was checked by western blot (see section below).

### Flow cytometry analysis

For the flow cytometry analysis, vSVZ single cell suspensions from mice at the specified ages were generated and stained for flow cytometry analysis as described (Llorens-Bobadilla *et al*, 2015). Briefly, vSVZ was microdissected using the Neural Tissue Dissociation kit with Trypsin and Gentle MACS dissociator (Miltenyi). Cells were stained with the following antibodies: O4 APC-Cy7 (1:100), GLAST (ACSA-1)-PE (1:50) (all from Miltenyi), Ter119 APC-Cy7 (Biolegend; 1:100), CD45 APC-Cy7 (BD; 1:200), Prominin1-APC (eBioscience; 1:75), Alexa488::EGF (Invitrogen; 1:100), and Sytox Blue (Invitrogen, 1:500). For analysis, we size selected the vSVZ cells and excluded for doublets, dead cells and CD45^+^/Ter119^+^/O4^+^ cells as recently described (Kalamakis *et al*, 2019). Signal acquirement was performed on a BD FACSCanto II at the DKFZ Flow Cytometry Facility and results were analysed using FlowJo v.10.

### Cell cycle analysis - CycleFlow

Cell cycle was assessed using CycleFlow as described (preprint: Jolly *et al*, 2020) with some modifications. CycleFlow was applied to NSCs with or without a pre-treatment of IFN-β. 24 h after cell seeding, either vehicle or IFN-β was added to NSCs for 17 h (this step was skipped in the non-pre-treated NSCs). Then, EdU from the Click-iT Plus EdU Flow Cytometry Assay Kit (ThermoFisher) was added at 10 µM. Cells were incubated 1 h at 37°C 25% CO_2_. Cells were collected, washed with DPBS and resuspended in pre-warmed medium containing either vehicle or IFN-β. Cells were then divided in different wells to be incubated and collected at the indicated incubation times (see Figure 1 and S1). At the collection time, cells were collected, dissociated (Accutase®, Sigma) and stained for viability with Zombie Red (Bioleged) following the manufacturer’s recommendations. After that, cells were washed and PFA-fixed using the Click-iT Plus EdU Flow Cytometry Assay Kit (ThermoFisher). Fixed cells were kept in 90% methanol DPBS at -20°C until completion of the time-course. Upon all collections, the Click-iT reaction was performed following the manufacturer’s recommendations in combination with a final staining with the DNA dye FxCycle Violet reagent (ThermoFisher; 1:1000). Signal acquirement was performed on a BD LSRFortessa Flow Cytometer and results were analysed using FlowJo v.10. Doublets and non-viable cells were excluded from the analysis. Doublets were discriminated using the DNA stain area vs. width. Cell-cycle progression mathematical inference was performed using uniform priors as already described (preprint: Jolly *et al*, 2020).

### Polysome profiling

NSCs were incubated with either vehicle or IFN-β as above explained. Right before cell collection, 100 µg/ml cycloheximide (CHX, EDM Millipore) in H_2_O was added to cells and they were incubated 5 min at 37°C 25% CO_2_. Cells were collected by centrifugation, washed with ice-cold DPBS containing 100 µg/ml cycloheximide (CHX), and lysed in the Polysome lysis buffer (20 mM Tris-HCl, pH 7.4, 120 mM KCl, 5 mM MgCl2, 14 mM β-mercaptoethanol, 1% NP-40, 100 µg/ml CHX, 1x complete protease inhibitor cocktail (Roche), 100 u/ml RNase Inhibitor (Thermo Fisher)). Lysates of differently treated NSC were loaded at equal amounts for total RNA onto preformed 17.5%-50% gradient of sucrose prepared in 20 mM Tris-HCl pH 7.4, 120 mM KCl, 5 mM MgCl2, 2 mM DTT, 100 µg/ml CHX. Free mRNPs, ribosome subunits, and polysome complexes were resolved by centrifugation for 2.5 h at 35,000 rpm in a SW41 rotor. The content of the tubes was fractionated with the Density Gradient Fractionation System (Teledyne ISCO) allowing for the recording of absorbance at 254 nm. 1 ml fractions were collected for RT-qPCR processing.

### RT-qPCR

Total RNA was extracted from cell cultures with the Arcturus PicoPure RNA Isolation Kit (Applied Biosystems). For the fractions of polysome profiling, RNA was purified via phenol/chloroform extraction as described (Faye *et al*, 2014). cDNA was synthesized with the SuperScript VILO cDNA synthesis kit (Thermo Fisher). DNA amplification and detection was performed with the PowerSYBR Green PCR MasterMix (Applied Biosystems) in C1000 Touch Thermal Cycler (Bio-Rad) using QuantiTect (Qiagen) primers (see Supplementary Table 2: RT-qPCR primers). For the calculation of relative gene expression, the ΔCt method was used. The distribution of analyzed mRNAs across polysome fractions was presented as a percentage of the total amount of the mRNAs for all fractions.

### Ribosomal footprinting

Cells were treated with vehicle or IFN-β 48 hours after seeding and incubated at 37°C 25% CO_2_ for the corresponding incubation time. Then, NSCs were incubated for 5 min with 0.1 mg/ml CHX at 37°C 25% CO_2_. Cells were collected, washed with DPBS supplemented with CHX, and lysed in 150 μl of the Polysome lysis buffer (see Polysome profiling section). Lysates containing 40 μg of total RNA were treated with 100 U of RNaseI (Ambion) for 45 min at room temperature, and the reaction was stopped by addition of 50 U of SUPERaseIn (Ambion). Generated 80S monosomes were collected via centrifugation through 25% sucrose cushion at 100,000 rpm for 1 h in a TLA100.2 rotor, and the pelleted RNA was extracted with the Direct-Zol RNA Miniprep kit (Zimo). Ribosomal RNA was depleted with the Gold Ribo-Zero kit (Illumina). Resulted RNA samples were resolved by electrophoresis in a 15% NOVEX TBE-Urea gel (Thermo Fisher). Gel slices including ribosome footprints corresponding to a fuzzy band of the 26-34 nt size range were cut out from the gel, crushed by centrifugation in 0.5 ml gel-breaker tubes (Segmatic) and extracted in 0.5 ml 10 mM Tris-HCl pH 7.0 by incubation for 15 min at 70°C. RNA was recovered by precipitation with isopropanol in presence of 0.3 M sodium acetate pH 5.5 and 20 μg Glycoblue (Thermo Scientific). Recovered RNA was dephosphorylated with T4 PNK (NEB) in the absence of ATP. Indexed libraries were generated using the SMARTer smRNA-Seq kit for Illumina (Takara).

For the parallel total mRNA sequencing, we used the same NSC lysates prepared as described above at the amount of 20 μg of total RNA. RNA was extracted with phenol/chloroform as described(Faye *et al*, 2014). RNA samples were treated with 3 U of TURBO DNase (Ambion) for 15 min at 37°C. RNA was again extracted with phenol/chloroform and recovered with ethanol precipitation in the presence of 0.3 m sodium acetate pH 5.2. Ribosomal RNA was depleted with the Gold Ribo-Zero kit (Illumina). After depletion 80 ng of each RNA sample was used to synthesized cDNA and make a library using the NEBNext Ultra Directional RNA library kit for Illumina (NEB).

Libraries quality was assessed using the Bioanalyzer 2100 (Agilent) and were sequenced in HiSeq 2000 v4.

For the analysis, Reads were trimmed applying the tool TrimGalore version 0.4.4_dev. The adaptor sequence “AGATCGGAAGAGC” (Illumina TruSeq, Sanger iPCR; auto-detected) as well as 3 bp from the 5’ end and 15 bp from the 3’ end were removed. In addition, sequences that became shorter than 18 bp (after quality trimming) were removed using TrimGalore’s default settings. After trimming reads had a length of 33 bp on average. Subsequently, reads were mapped to the mm10 transcriptome build GRC38.93 from ENSEMBL using boWTie version 0.12.7 with its standard options. Reads falling into genes were counted from BAM files applying a suited R/Bioconductor workflow (function SummarizeOverlaps with mode ‘Union’). Duplicated reads were removed. For ribosomal footprinting samples, reads in the whole gene body or in the coding part of genes (CDS) were counted separately, for total RNA samples reads in the whole gene body were counted. Next, to get an estimation of the translation efficiency (TE) per gene, log-fold-changes between ribosome protected reads from the CDS and total RNA samples were computed applying DESeq2. DESeq2 was chosen as it is considered a standard tool for modelling negative-binomial distributions arising in sequencing experiments. The likelihood ratio test (LRT) was applied on the TE analysis upon IFN-β treatment. Genes with positive fold-changes are considered to be enhanced, those with negative ones to be repressed. The gene set enrichment analysis was performed using the clusterProfiler tool from Bioconductor using fold-changes for all expressed genes. Gene ontology terms related to “biological process” with an FDR < 0.05 and highest significance were selected for the plots.

### Proteome and phosphoproteome

Cells were treated with vehicle or IFN-β and incubated at 37°C 25% CO_2_ for the time specified in the figures. Cell lysates for the proteome and phosphoproteome analysis were prepared as described (Potel *et al*, 2018) with some modifications. Briefly, cells were collected, washed and weighted to determine the mass of cells pellets. One volume of the cell pellet was resuspended in six volumes of proteome lysis buffer composed of 100 mM Tris-HCl pH 8.5, 7 M Urea, 1 mM MgCl_2_, 1 mM sodium ortovanadate, 1% Trition X-100, 1X PhosphoSTOP inhibitor (Roche), 1X Complete EDTA free protease inhibitor (Roche). To improve lysis, cells suspension was sonicated 3 times for 10 s (1 sec on, 1 sec off) with 30 s pause at 40% output using the Fisherbrand Model 120 Sonic Dismembrator (Fisher Scientific). The lysate was clarified by centrifugation at 21000 g, for 1 h at 4°C. Lysates were incubated for 2 h at room temperature. Protein concentration was determined with the Pierce BCA Protein Assay Kit (Thermo Fisher) and 400 µg of total protein for each sample were subjected to metal-affinity enrichment & mass spectrometry at the DKFZ Genomics and Proteomics Facility. Proteins and phosphopeptides were quantified and analyzed for differential abundance using Perseus45.

### Western blot analysis

Cells were collected, washed with DPBS and resuspended in 100 µl of 0.2% SDS, 1 mM vanadate, 1X cOmplete Protease inhibitor cocktail (Roche), 1X PhosphoSTOP inhibitor (Roche) and pass several times through a 23G syringe needle. 15 µg of total protein from cellular lysates were resolved in NuPAGE 10% gels (Thermo Fisher) and transferred to nitrocellulose membranes. The membranes were blocked in 5% BSA in TBS-T buffer and probed overnight with primary antibody. Membranes were washed and reprobed with secondary anti-rabbit or anti-mouse HRP-antibody (Dianova). The signals were developed with the Western Lightning Plus ECL substrate (Perkin Elmer) and scanned using the ChemiDoc Touch Imageing System (Bio-Rad). Loading controls were always run on the same blot than the quatified phospho- or total protein. The Image Lab version 5.2 (Bio-Rad) was used for band intensity quantification.

### Luciferase assay

For nucleofection, 3 x 10^6^ NSCs were mixed with 5 µg of corresponding DNA with the Amaxa P4 Primary Cell 4D-Nucleofector X Kit S (Lonza) and pulse was delivered using the CA137 programme in a 4D-Nucleofector X Unit (Lonza). Afterwards, the cells were washed, resuspended in medium and separated in three technical replicates. After recovery, cells were treated with IFN-β, Torin1, a combination of both or their corresponding vehicles as stated in the figures. Then, cells were washed and lysed in 1x Passive Lysis buffer from the Dual-Luciferase Reporter Assay System (Promega). Lysis was proceeded on an orbital shaker for 15 min at room temperature. Luciferase Assay reagent was added to each sample, mixed and all volume was transferred into a 96 well White Cliniplate (Thermo Fisher). For *Firefly* luciferase activity the plate was scanned in a Synergy LX mutli-mode reader (BioTek). Then 100 µl of Stop&Glo reagent was added and the plate was scanned again for *Renilla* luciferase. *Renilla* luminescence was normalized to *Firefly* luminescence (with technical triplicates) and the final results were presented as fold change to the control samples.

### Plasmid construction

DNA constructs with different WT and mutated 5’UTRs of mouse *Sox2* mRNA were assembled in the psiCheck-2 vector (Promega) including synthetic *Renilla* luciferase gene (hRluc) driven by SV40 early enhancer/promoter and synthetic *Firefly* luciferase gene (hluc+) under HSV-TK promoter. The UTRs flank hRluc open reading frame. The full-length 5’UTRs of mouse *Sox2* (NM_011443.4) and *Rps21* (variant2, NM_025587.2) as well as their mutated variants carrying the deletions of PRM in *Sox2* (5UTRmut *Sox2*; CTCTT deleted), PRM in *Rps21* (5UTRmut *Rps21*; TCCTTTC deleted) were synthesized and inserted into pEX-K4 (*Sox2*) or pEX-A2 (*Rps21*) by Eurofins. The plasmids were used to amplify the UTRs with Phusion High-Fidelity DNA polymerase (NEB) using corresponding primers carrying SfiI and NheI restriction sites to allow the inserts between SV40 promoter and the reading frame of hRluc. The used primers are listed in the Supplementary Table 1. The final plasmids were designated as 5Sox, 5mutSox, 5Rps21, 5mutRps21.

The full-length 5’UTR of mouse beta actin (*Actb*) mRNA (NM_007393.5) was generated by amplification of cDNA library prepared from NSCs isolated from C57BL/6 mice. For generation of the 5’UTR *Actb*, the used PCR primers were flanked with SfiI and NheI sites to allow the insertion in front of hRluc reading frame in psiCheck-2 vector (Promega).

The accuracy of cloning was verified by sequencing of the inserts in all generated plasmids from both directions with corresponding primers for sequencing (see Appendix Table 3).

### Single-cell transcriptomics

To characterize the single-cell transcriptomics of NSCs and their progeny, we made use of TiCY (referred to as IFNAGR^WT^) and TiCY-IFNAGR KO mice (referred to as IFNAGR^KO^). In these mice, tamoxifen-induced Cre recombination takes place in neural stem cells in the vSVZ, which express *Tlx* (*Nr2e1*) (Liu *et al*, 2008), and will stably activate the production of eYFP labelling NSCs and their progeny.

Tamoxifen (TAM) injection was done in three days as follows: two doses daily of 1mg of tamoxifen in 100 uL of a solution of 10 mg/ml of TAM in EtOH 10% diluted on sunflower seed oil. TiCY-IFNAGR^WT^ young mice were injected with TAM at 10 weeks old and were sacrificed 6 weeks afterwards. TiCY-IFNAGR^WT^ old mice (71 weeks old), TiCY-IFNAGR^KO^ young (7 weeks old) and old mice (85 and 98 weeks old) were injected with TAM and were sacrificed 9 weeks afterwards. After animal sacrifice, the vSVZ, striatum, rostral migratory stream and olfactory bulb was isolated. These latter three tissues were pooled together and referred to as Rest of the Brain (RoB). Tissues were processed as described previously (Kremer *et al*, 2021) and sorted in a BD FACSAria II at the DKFZ Flow Cytometry Facility. For sorting, we size-selected the vSVZ or RoB cells and excluded for doublets, dead cells and CD45^+^/Ter119^+^/O4^+^ cells as recently described (Kalamakis *et al*, 2019). We sorted vSVZ eYFP^+^ cells and GLAST^+^ cells. From the RoB, cells that were eYFP^+^ and eYFP^-^/PSANCAM^low^ were sorted. For every condition, 2 mice were used whose cells were labelled with Hashtags oligos (HTO) using the Cell Hashing protocol from Biolegend (TotalSeq-A). All the sorted cells were pooled on the same tube and processed in the Chromium Next GEM Chip G using Chromium Next GEM Single Cell 3’ v3.1.

Gene expression libraries were prepared following the manufacturer’s protocol (Chromium Next GEM Single Cell 3’ v3.1) at the Single-cell open lab at DKFZ and sequenced on a NextSeq2000 V3 PE 100bp at the Sequencing open lab provided by the DKFZ Genomics and Proteomics Core Facility. Each condition (genotype and age combination) was processed on a separate 10X reaction. Hashtags libraries were processed according to the manufacturer’s instruction (Biolegend) and were sequenced on a NextSeq 500 Mid output PE 75bp at the DKFZ Genomics and Proteomics Core Facility.

### Single-cell transcriptomic data analysis

To derive a “NSC type I Interferon Response” signature we used the 16h IFN-β treated Ribo-Seq transcriptomic libraries. Reads were trimmed using trimgalore 0.6.6 using default settings and mapped and quantified using STAR 2.7.7a with --quantMode GeneCounts. Star outputs were concatenated and read into R 4.0.5 where DESeq2 1.30.1 was used to compute differential expression between treated and untreated samples. The genes with the 300 most significantly changed genes after multiple-testing correction from a one-sided Wald test (increased expression) were chosen as our “NSC Type I Interferon Response” gene set.

For the single-cell libraries analysis, we filtered them using the 10x index-hopping-filter 1.0.1 and thereafter quantified using kallisto|bustools’s lamanno workflow (kb_python 0.26.3). Hashtag libraries were quantified using CITE-seq-Count 1.4.5. Count matrices were read into scanpy 1.6.0 and preprocessed by filtering out genes present in less than three cells and cells with less than 100 genes. Doublets were called and removed with scrublet 0.2.1. Cells with less than 10% mitochondrial UMIs, 70-6000 genes and 1000-20000 UMIs were retained as quality cells. Counts were normalised (to a target sum of 10000) and log1p transformed using scanpy methods. A UMAP was computed from 50PCs computed on scaled counts. Main clusters were identified in UMAP using DBSCAN (scikit-learn 0.22.2.post1). An auxilliary leiden clustering was computed. The largest DBSCAN cluster was identified as the lineage cluster based on marker expression. Non-subventricular astrocytes were identified and removed from this cluster by inspecting the fraction of SVZ hashtags in each cell. Other clusters were named by summed expression of celltype specific-markers (Pecam1, Prom1 and Cldn5 for endothelials, Itgam and Ptprc for microglia, Rbfox3, Calb2, Nefl and Th for neurons). Diffusion pseudotime (from scanpy) was run on the lineage cluster using the cell with the highest Aqp4 expression as the root cell. Pseudotime was then binned into qNSC1, qNSC2, aNSC, TAP and NB based on visual inspection of the expression of well-known markers Aqp4, Egfr, Dcx, Mki67, S100b. Hashtag samples were called for each cell as the maximum posterior from a multinomial mixture model (R 4.0.5 package mixtools 1.2.0) with 5 components (one for each hashtag and an extra component for no hashtags).

Gene set scores were then computed for each cell by summing up the normalised (per cell to 1000000 UMIs) and log1p-transformed expression of genes from a given set. Group-wise gene set scores were then computed as the avearge of all of the cell’s scores in a given group of interest. Scores were computed for each replicate, celltype, age and genotype combination for MSigDB’s “HALLMARK_INFLAMMATORY_RESPONSE”, “HALLMARK_INTERFERON_ALPHA_RESPONSE” genesets, as well as the gene set identified by Wu et al. 2018 (Wu *et al*, 2018) and our “NSC Type I Interferon Response”.

### IFNAGR KO Modelling

We built on the previously established mathematical model of cell population dynamics (Kalamakis *et al*, 2019), describing counts of quiescent (non-cycling) and active (cycling) stem cells. The model is given by a system of ordinary differential equations:

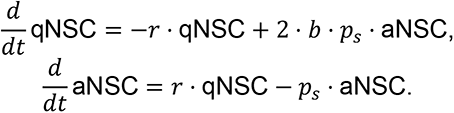

The parameter *r* describes the activation rate that may exponentially decrease in time

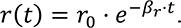

Self-renewal is modelled by a function *b*

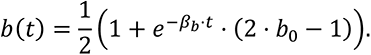

The choice of this function follows the assumptions that self-renewal probability is not larger than ½ and may be a constant or increasing in time function.

The remaining parameter *p_s_* = log(2)/(17.5/24) ≈ 0.9506 corresponds to the cell cycle rate. It was chosen from literature, following our previous publication (Kalamakis *et al*, 2019). In summary, the model is characterised by free parameters *r_0_*, *b_0_*, *β_b_* and *β_r_*. The initial value additionally provides the parameter NSC_0_ which was used to compute initial values for the specific compartments from the steady state ratio (Appendix Data 1).

Parameters were estimated using a multi-start box-constrained weighted-least-squares approach. The objective functions were built using new FACS quantifications for the total numbers of stem cells and the fraction of actively cycling neural stem cells from our previous publication. We applied solvers provided and chosen by the DifferentialEquations.jl package. Weights were computed as the inverse standard deviation for each age and genotype combination. Starting values were sampled from within reasonable bounds using Latin Hypercubes computed by the LatinHypercubeSampling.jl library, each was then optimized using a box constrained optimizer from the Optim.jl library. The best parameter estimates were kept as the optimum. To aid in model selection, AIC*_c_* was computed for each model and dataset combination.

Finally, we estimated parameters for models with population-dependent self-renewal probabilities, described by a Hill function:

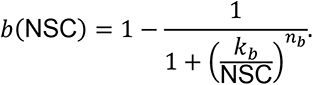

We estimated these with a single *k_b_* for both genotypes. The parameter *n_b_* was estimated individually per-genotype (IFN-dependent self-renewal) or shared (IFN-independent self-renewal) to provide the two scenarios of self-renewal dependence on interferons.

### IFNAGR KO simulations at arbitrary time points

To simulate the age-specific dynamics of the IFNAGR KO, we built a model with a time-dependent switch in the parameter functions. At the intervention age, the parameter functions are switched from the previously estimated WT parameter functions to their IFNAGR KO counterparts. We simulated this intervention model for varying intervention ages. For each simulation we computed the number of stem cell loss at 660 days age compared to the wildtype. We also computed the progenitor production as the cell flux from active neural stem cells to progenitors. We then integrated progenitor production from 0 to 700 as the total life-long progenitors produced.

### Statistical analysis

Biological replicates “n” in the figures refer to biological replicates as of either different mice or different NSCs cultures extracted from different mice. Plotted bars represent mean and plotted error bars represent standard deviation from different biological replicates. Only biological replicates were considered for statistical analyses. Statistical tests were performed as indicated in each figure legend with a significance level of α = 0,05. To test for biphasic response in western blot data, linear models explaining log2 fold changes were fitted with a single explanatory variable of time being greater than a change time. To find this change time, models were fitted for varying change times and the model with the lowest AIC was chosen. To determine the significance of this biphasic response, one-way ANOVA was performed on this model.

### Data and code availability

Raw Single-cell RNA-seq and Ribo-Seq data have been deposited at the Gene Expression Omnibus (GEO) and are publicly available with GEO Accession Number GSE197217 as of the date of publication. All original code will be deposited at https://github.com/Martin-Villalba-lab/Data/ and publicly available as of the date of publication.

Any additional information required to reanalyse the data reported in this paper is available from the lead contact upon request.

### Schematic illustrations

Schemes were created with BioRender.com and Adobe Illustrator.

## Acknowledgments

We thank S. Wolf & D. Helm from the DKFZ Genomics and Proteomics Core Facility; S. Schmitt from the DKFZ Flow Cytometry Core Facility; J.-P. Mallm from the DKFZ scOpenLab; the DKFZ Centre for Preclinical Research; K. Volk, K. Menzner & H. Abendroth for technical assistance; B. Bukau and the members of the Martin-Villalba laboratory for critical comments.

This work was supported by the European Research Council (ERC; REBUILD_CNS), the Helmholtz Zukunftsthema ageing and Metabolic Programming (AMPro) ZT-0026, the German Research Foundation (DFG; SFB873), and the German Cancer Research Center (DKFZ).

## Author contributions

**Damian Carvajal Ibanez**: Conceptualization, Formal Analysis, Investigation, Validation, Methodology, Project administration, Resources, Visualization, Writing – original draft, Writing – review & editing. **Maxim Skabkin**: Conceptualization, Formal Analysis, Investigation, Methodology, Project administration, Resources, Visualization, Writing – review & editing. **Jooa Hooli**: Conceptualization, Formal Analysis, Data curation, Software, Validation, Resources, Visualization, Writing – original draft, Writing – review & editing. **Santiago Cerrizuela**: Investigation, Methodology, Resources, Writing – review & editing. **Manuel Goepferich**: Formal Analysis, Data curation, Software, Resources, Visualization. **Adrien Jolly** and **Matilde Bertolini**: Resources. **Marc Zumwinkel**: Investigation. **Thomas Hoefer**, **Guenter Kramer**, **Simon Anders** and **Aurelio Teleman**: Supervision, Resources. **Anna Marciniak-Czochra**: Conceptualization, Project administration, Supervision, Writing – original draft, Writing – review & editing. **Ana Martin-Villalba**: Conceptualization, Project administration, Supervision, Funding acquisition, Writing – original draft, Writing – review & editing.

---

**Expanded View Figure 1.**
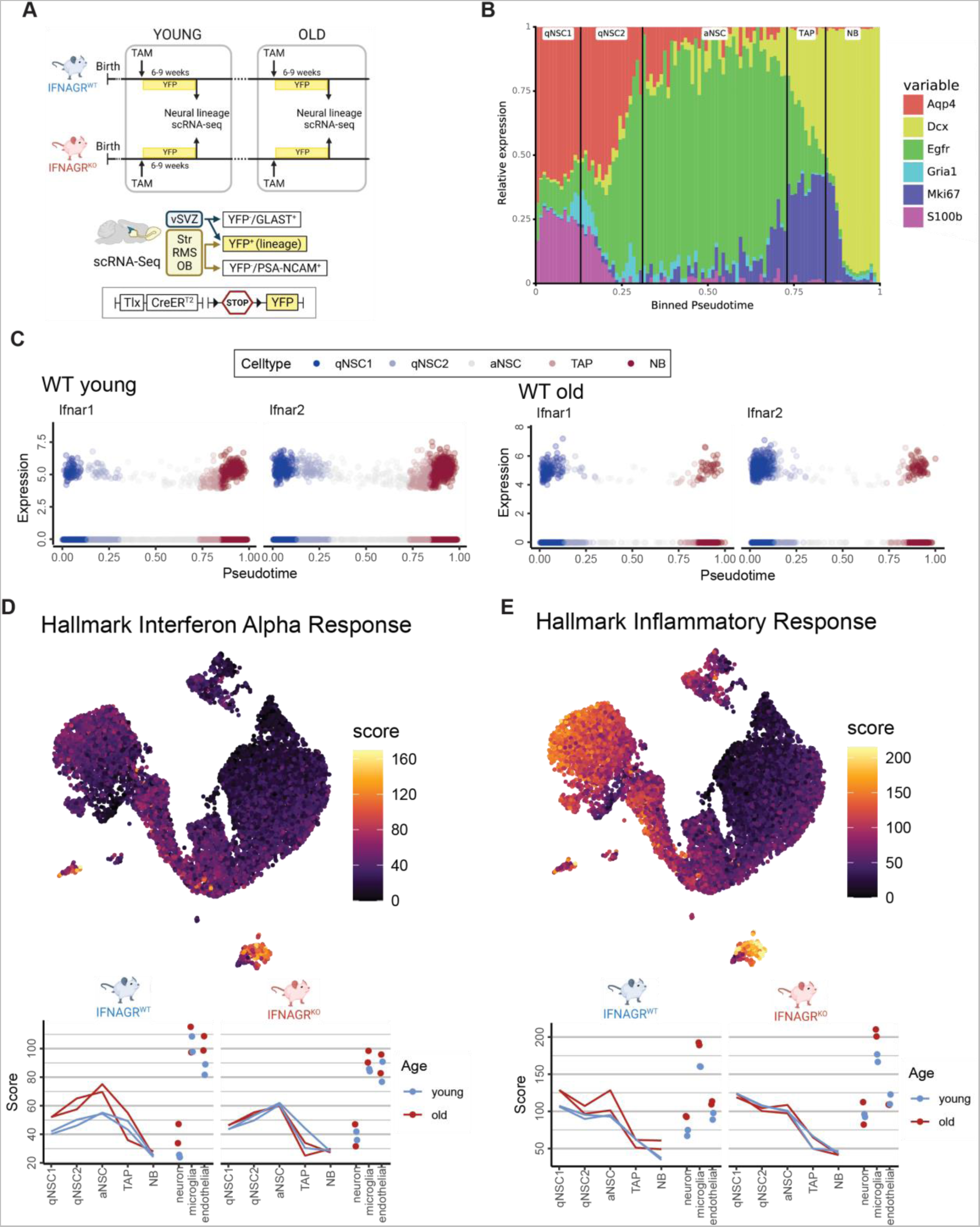
Inference of IFN signatures in the vSVZ niche. **A.** Experimental layout for scRNA-Seq in young and old mice lacking interferon receptors. All mice represent TiCY (Tlx reporter, see methods and Fig EV1) that are either IFNAGR WT or KO. vSVZ (Ventricular Subventricular Zone), RMS (Rostral Migratory Stream), OB (Olfactory Bulb), TAM (Tamoxifen). **B.** Relative gene expression of relevant markers for cell types along pseudotime. Black lines denote cuts between cell types. **C.** Relative expression of type-I IFN receptors in young and old wildtype cells over pseudotime colored by lineage cell types. **D.** Scores computed for the Hallmark Interferon Alpha Response signature displayed in the UMAP embedding for young cells (with colours clipped to the range seen in the lineage cells) and averaged for the cell types in our analysis at varying ages in in IFNAGR^WT^ and IFNAGR^KO^ cells. n = 2 biological replicates per age and genotype. **E.** Scores computed for the Hallmark Inflammatory Response signature displayed in the UMAP embedding for young cells (with colours clipped to the range seen in the lineage cells) and averaged for the cell types in our analysis at varying ages in IFNAGR^WT^ and IFNAGR^KO^ cells. n = 2 biological replicates per age and genotype.

**Expanded View Figure 2.**
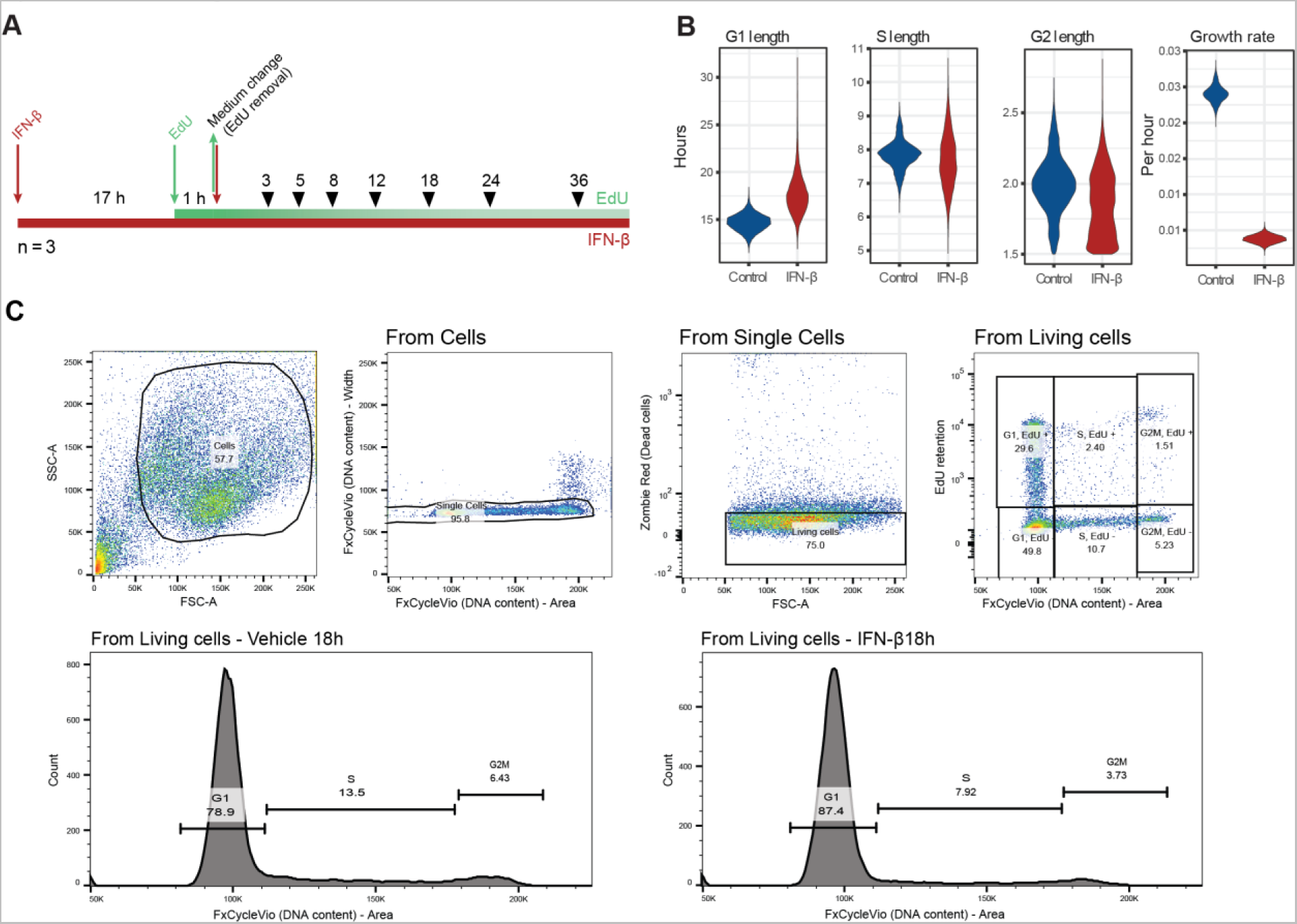
Cycleflow anaylsis: IFN-β induce cell cycle exit in NSCs. **A.** EdU and IFN-β exposure experimental scheme for Cycleflow. **B.** Cell cycle properties inferred from Cycleflow in IFN-β-treated NSCs. n = 3 biological replicates. **C.** Gating strategy for quantification of cell cycle states for Cycleflow or general cell cycle stages G_1_, S and G_2_M (lower panels).

**Expanded View Figure 3.**
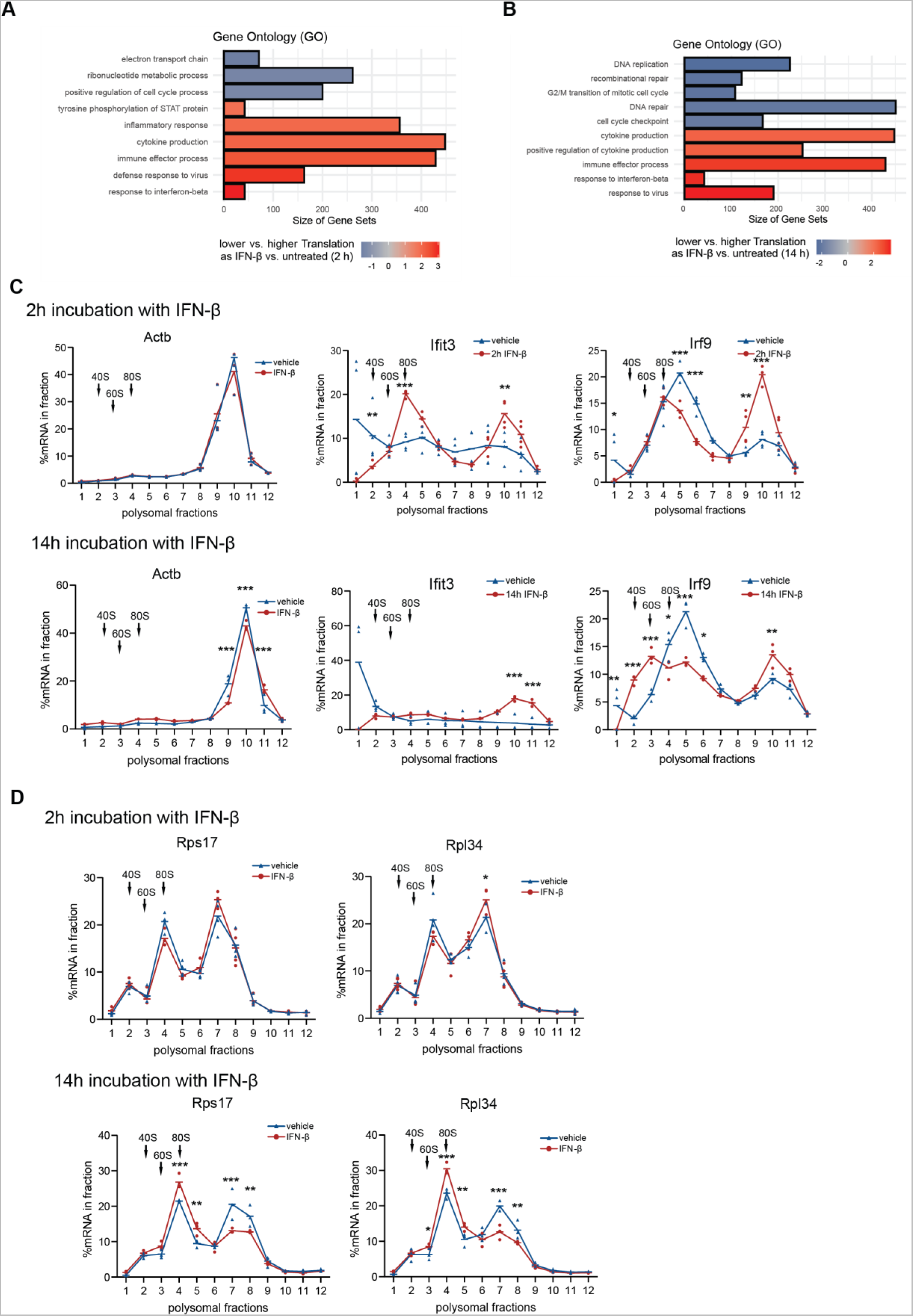
IFN-β induce a biphasic control of mRNA translation in NSCs. **A,B.** Results of Ribo-Seq depicting translation efficiency as the interaction of log2 fold changes (LFC) between footprints (ribosome protected reads) and total RNA at 2h and 14h IFN-β treatment. FDR < 10%, LR-Test. Genes with a p-value <0.1 after FDR correction are highlighted. The associated GO terms are depicted. n = 3 biological replicates. **C.** Polysome profiling (RT-qPCR) of actin beta and of the interferon stimulated genes (ISGs) Ifit3 and Irf9 upon 2h and 14h IFN-β treatment. Hyphens represent mean of biological replicates. Arrows indicate the 40S, 60S and 80S subunits of the ribosome. n = 3-4 biological replicates. **D.** Polysome profiling (RT-qPCR) of *Rps17* and *Rpl34* upon 2h and 14h IFN-β treatment. Hyphens represent mean of biological replicates. Arrows indicate the 40S, 60S and 80S subunits of the ribosome. n = 3-4 biological replicates. Two-way ANOVA with Šídák’s multiple comparison test was computed. Outliers of fraction 1 from Ifit3 were excluded from the statistical analysis. *p<0.05, **p<0.01, ***p<0.001.

**Expanded View Figure 4.**
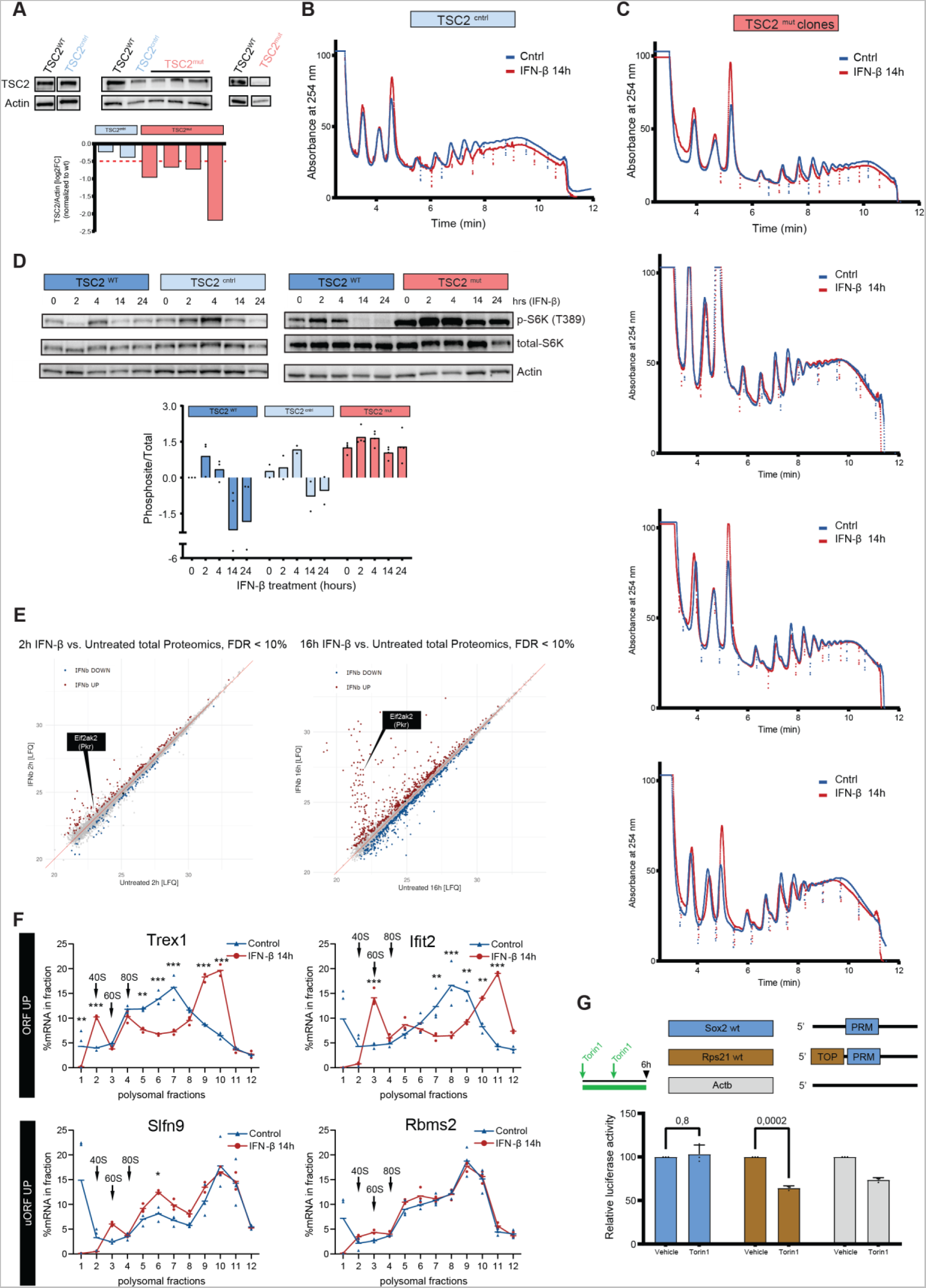
The biphasic control of mTORC1 by IFN-β relies on TSC2, PKR and eIF2α. **A.** WB images and quantifications of TSC2 relative to actin beta and normalized to TSC2^WT^ NSCs in different CRISPR-mutated (TSC2^mut^) or CRSPR-non-mutated (TSC2^cntrl^) NSC clones. **B,C.** Polysome profiles of CRISPR-mediated TSC2^cntrl^ and TSC2^mut^ NSC clones treated with IFN-β (red) as compared to control (blue). Each plot represents different TSC2^mut^ NSC clones. **D.** Representative WB images and quantifications (log2FC) of p-p70S6K^Thr389^ in TSC2^WT^, TSC2^cntrl^ & TSC2^mut^ NSCs treated with IFN-β and normalized to vehicle (t=0h) TSC2^WT^. Bars represent the mean value. n = 2-4 clonal replicates. **E.** Relate label-free quantification (LFQ) of proteomics from WT NSCs untreated or treated with IFN-β for 2h or 16h. n = 5 biological replicates. **F.** Polysome profiling (RT-qPCR) of genes displaying upregulation of footprints in the ORF (Trex1 and Ifit2) compared to genes with an upregulation in uORFs (Slfn9 and Rbms2) upon 14h IFN-β treatment. Hyphens represent mean of biological replicates. Arrows indicate the 40S, 60S and 80S subunits of the ribosome. n = 3 biological replicates. Two-way ANOVA with Šídák’s multiple comparison test was computed. Outliers of fraction 1 from Ifit2, Slfn9 and Rbms2 were excluded from the statistical analysis. **G.** Experimental layout (left) & 5’UTR constructs priming *renilla* luciferase. TOP = 5’Terminal Oligopyrimidine motif; PRM = 5’ Pyrimidine Rich Motif. Luciferase activity (right) controlled by the upstream 5’UTR fragment from Sox2, Rps21 and Actb. Data is normalized to vehicle and is represented as mean ± SD. n = 3 biological replicates. Two-way ANOVA test with Ŝídák’s multiple comparison test (p-value specified). *p<0.05, **p<0.01, ***p<0.001.

**Expanded View Figure 5.**
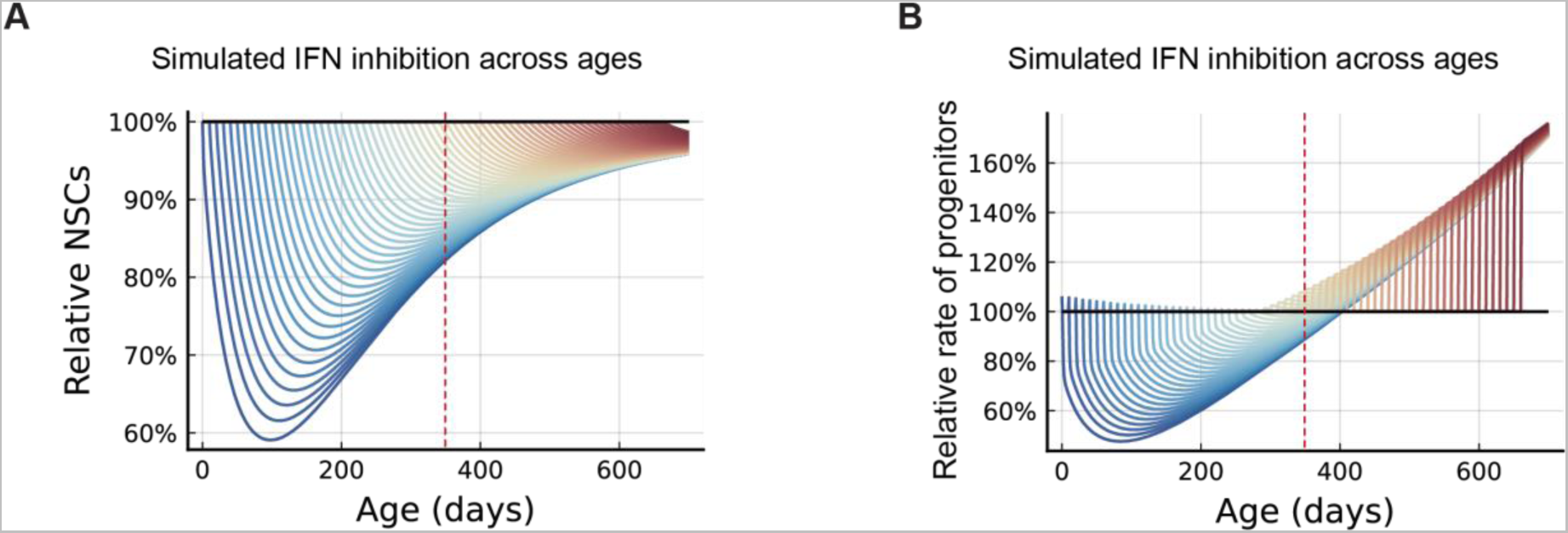
Modelling of interferon receptors removal along lifespan. **A.** Induced interferon knockout simulations (colored, each color represents a different intervention timepoint) of relative loss of stem cells across age compared to wildtype simulations (black; 100%). Dashed red line denotes age 350 days. **B.** The relative rate at which progenitors are produced (2*p_s_* (1 – *b*) *aNSC*) from NSCs for induced interferon knockout simulations with interferon dependent self-renewal (colored, each color represents a different intervention timepoint) compared to WT simulations (black; 100%). Dashed red line denotes age 350 days.

**Appendix Table 1:** Phosphosites

| Protein | Phosphosite | log2FC (2h) | log2FC (16h) |
| --- | --- | --- | --- |
| Akt1 | S473 | 0.1 | -1 |
| Irs1 | Y608 | 1.2 | 0.3 |
| Irs1 | S1134 | 0.9 | -2.3 |
| Irs1 | S1137 | 0.2 | -1.7 |
| Eif2ak2 | S32 | 0.1 | 4.2 |
| Eif2ak2 | S110 | 0.4 | 3.3 |
| Eif2ak2 | S163 | -0.2 | 2.4 |
| Eif2ak2 | S223 | 0.5 | 2.1 |
| Eif2ak2 | S240 | -1 | 3.1 |
| Ccnd1 | T286 | 1.9 | 0.3 |

